# Integrating ecosystem and contaminant models to predict the effects of ecosystem fluxes on contaminant dynamics

**DOI:** 10.1101/2023.07.15.549171

**Authors:** Anne M. McLeod, Shawn J. Leroux, Matteo Rizzuto, Mathew A. Leibold, Luis Schiesari

## Abstract

Pollution is one of the major drivers of ecosystem change in the Anthropocene. Toxic chemicals are not constrained to their source of origin as they cross ecosystem boundaries via biotic (e.g., animal migration) and abiotic (e.g., water flow) vectors. Meta-ecology has led to important insights on how spatial flows or subsidies of matter across ecosystem boundaries can have broad impacts on local and regional ecosystem dynamics but has not yet addressed the dynamics of pollutants. Understanding how these meta-ecosystem processes on contaminant dynamics may reverberate up a food chain is important even if they might be difficult to predict. Here we derive a modelling framework to predict how spatial ecosystem fluxes can influence contaminant dynamics and how the severity of this impact is dependent on the type of ecosystem flux leading to the spatial coupling (e.g., herbivore movement vs abiotic chemical flows). We mix an analytical and numerical approach to analyze our integrative model which couples two distinct sub-components – an ecosystem model and a contaminant model. We observe an array of dynamics for how chemical concentrations change with increasing nutrient input and loss rate across trophic levels. When we tailor our range of chemical parameter values to specific organic chemicals our results demonstrate that increasing nutrient input rates can lead to trophic dilution in pollutants such as polychlorinated biphenyls across trophic levels. Yet, increasing nutrient loss rate causes an increase in concentrations of chemicals across all trophic levels. A sensitivity analysis demonstrates that nutrient recycling is an important ecosystem process impacting contaminant concentrations, generating predictions to be addressed by future empirical studies. Importantly, our model demonstrates the utility of our framework for identifying drivers of contaminant dynamics in connected ecosystems including the importance that a) ecosystem processes, and b) movement, especially movement of lower trophic levels, have on contaminant concentrations. For example, how increasing nutrient loss rate leads to increasing contaminant concentrations, or how movement of lower trophic levels contributes to elevated herbivore contaminant concentrations. This dynamic is particularly relevant given that the flow of matter between ecosystems also serves as a vector for the transport of contaminants.

## INTRODUCTION

Ecosystems are coupled in space through the movement of energy, materials, and organisms (Loreau et al. 2003, Marleau et al. 2020). This movement can have strong impacts on local dynamics, altering the spatial distribution of resources (Gravel et al. 2010), rescuing local populations from extirpation (Hanski 1998), and triggering trophic cascades (Polis et al. 1997, Leroux and Loreau 2008). Local effects of the flow of energy, material and organisms across ecosystems (i.e., subsidies) can also have broad impacts at regional scales (Harvey et al. 2020), for example, across watersheds (e.g., Harvey et al. 2017, McCann et al. 2021) and coastal areas (Menge et al. 2015). In recent years there has been a proliferation of both theoretical (see Loreau et al. 2003, Massol et al. 2011, Gounand et al. 2018) and empirical (see reviews in Allen and Wesner 2016, Montagano et al. 2019) studies exploring the local consequences of spatial connections and the impacts of environmental change on ecosystem processes (Larsen et al. 2016). This work has focused primarily on resource subsidies, but the flows of materials and organisms can also serve as vectors for the movement of other substances including contaminants—with unsuspected, and often overlooked consequences (e.g., Blais et al. 2007, Schiesari et al. 2018, Kraus et al. 2020).

Anthropogenic contaminants are now ubiquitous (Malaj et al. 2014, Walters et al. 2016). In fact, pollution is one of nine anthropogenic activities threatening to push the Earth beyond the unusually stable state that the planet has been in for the past 10 000 years (Rockström et al. 2009), one of the five most important direct drivers of ecosystem change (Nelson 2005), and a leading cause of extinctions (e.g. Wilcove et al. 1998). Moreover, chemical contamination is not necessarily an isolated stressor—instead, increased contaminant inputs are often coupled with other common stressors such as elevated nutrients and invasive species (Burton Jr et al. 2017).

Despite evidence of a concurrent rate of increase in the production of chemical contaminants with that of global fertilizer use (Bernhardt et al. 2017), the effects of chemical contaminants are often studied in isolation from ecosystem studies examining the effects of ecological subsidies, such as nutrients (Bernhardt et al. 2017, Schiesari et al. 2018). Given the rapid changes in biotic and abiotic processes in the Anthropocene, understanding how ecosystem context and ecosystem change interact with contaminant dynamics is critical for predicting contaminant exposure and its consequences. While there have been several studies that present conceptual models describing feedbacks between ecosystems and contaminants (e.g., see discussion in Schiesari et al. 2018; and Chumchal and Drenner 2020, Kraus et al. 2020), we lack a more precise mathematical framework coupling conventional ecosystem processes with contaminant dynamics (see Figure 1) and hence we also lack an understanding of how variation in spatial ecosystem processes may influence contaminant dynamics across ecosystems (see discussion in Muehlbauer et al. 2020).

**Figure 1.**
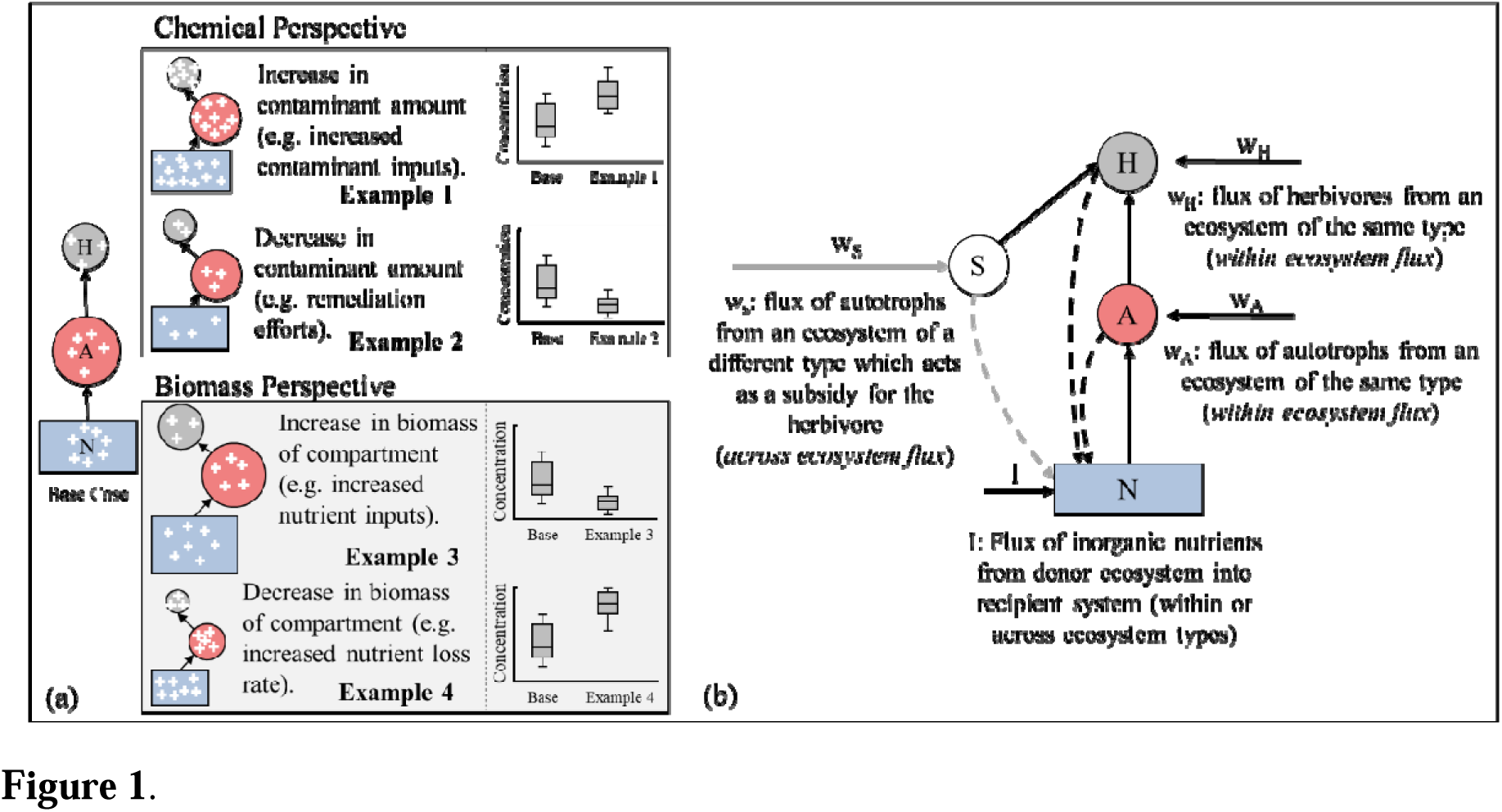
(a) Contaminant concentration can be altered in three ways; (i) change in the mass of the chemical in the compartment (denoted by white crosses; example 1 and 2) and explored in Appendix S5, (ii) change in the biomass of the compartment (demonstrated by change in the size of the circles and/or square where circles represent the entire biomass of that compartment; example 3 and 4), or (iii) change in both the mass of chemical in the compartment and change in the biomass of the compartment. Here, we focus on (ii) or how spatial *ecosystem processes* (e.g., the movement of organisms which transport nutrients and contaminants to the recipient ecosystem, or flooding events which lead to an increase in nutrient inputs in the recipient ecosystem) impact contaminant concentrations. In this simple example, we can see that two very different mechanisms can give rise to similar patterns (e.g., example 1 and example 4 both demonstrate elevated contaminant concentrations). It is only by coupling a contaminant model with an ecosystem model into a single modeling framework that we can tease out the individual and combined effects of stressors and responses. Here, we use an ecosystem model (b): a model that explicitly includes biotic and abiotic components and interactions among these components. We use this model to examine how fluxes of biotic and abiotic (*I*) material, energy, or organisms across ecosystems influence contaminant concentrations in a recipient ecosystem (*w_H_*, *w_A_*, and *w_S_* where *w_H_*, and *w_A_* are within ecosystem fluxes and *w_S_* is an across ecosystem flux). See Appendix S4: Figure S1 for full diagram of the model and Table 1 for model equations, and variable and parameter definitions.

**Table 1.**
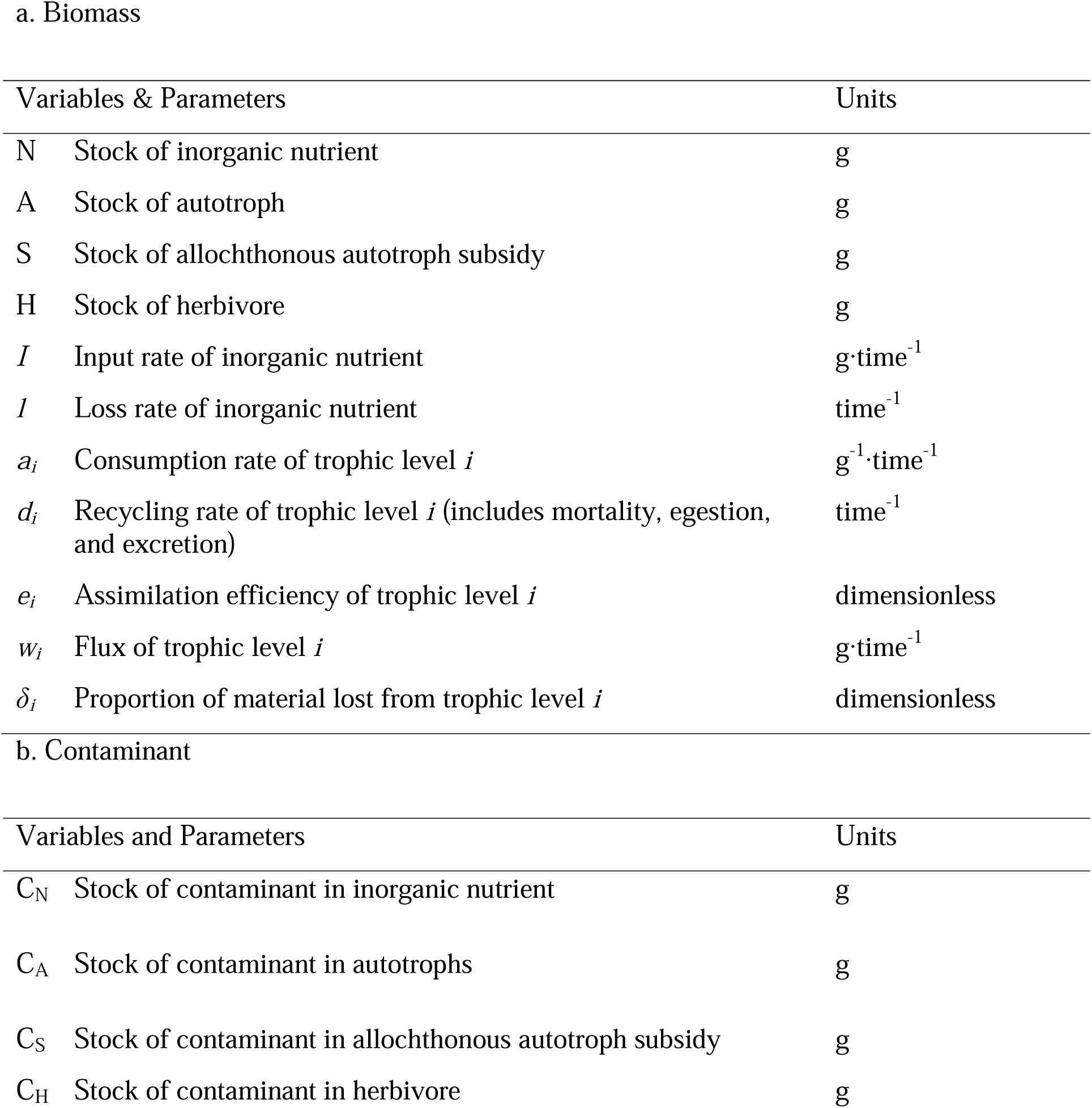
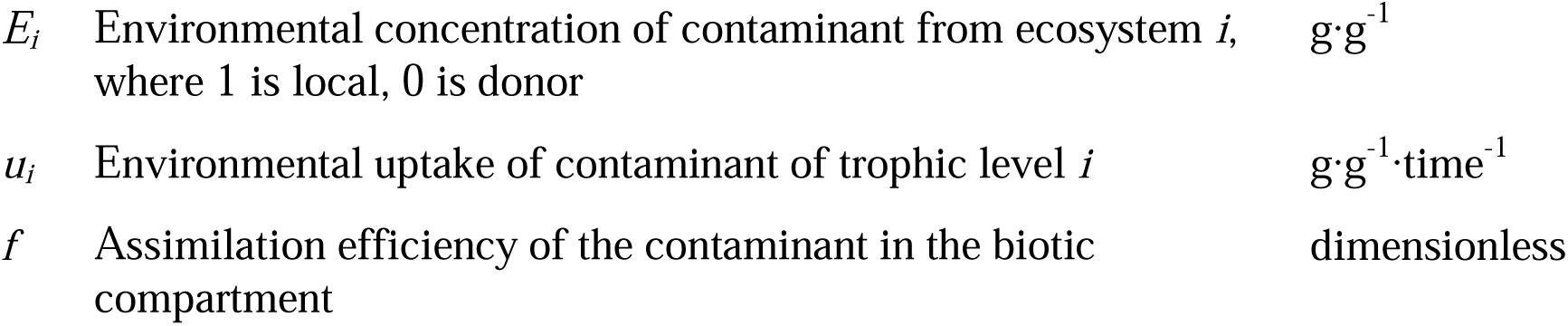
Description of full model equations and variables and parameters definitions with units for (a) biomass and (b) contaminant. Individual models presented in Appendix S4: Figure S1, can be recovered from the full model equations listed here. For example, the base model is recovered by setting *w*_A_ = *w*_S_ = *w*_H_ = 0.

| Variables & Parameters | Units |
| --- | --- |
| N Stock of inorganic nutrient | g |
| A Stock of autotroph | g |
| S Stock of allochthonous autotroph subsidy | g |
| H Stock of herbivore | g |
| $I$ Input rate of inorganic nutrient | $\text{g} \cdot \text{time}^{-1}$ |
| $l$ Loss rate of inorganic nutrient | $\text{time}^{-1}$ |
| $a_i$ Consumption rate of trophic level $i$ | $\text{g}^{-1} \cdot \text{time}^{-1}$ |
| $d_i$ Recycling rate of trophic level $i$ (includes mortality, egestion, and excretion) | $\text{time}^{-1}$ |
| $e_i$ Assimilation efficiency of trophic level $i$ | dimensionless |
| $w_i$ Flux of trophic level $i$ | $\text{g} \cdot \text{time}^{-1}$ |
| $\delta_i$ Proportion of material lost from trophic level $i$ | dimensionless |

b. Contaminant
| Variables and Parameters | Units |
| --- | --- |
| $C_N$ Stock of contaminant in inorganic nutrient | g |
| $C_A$ Stock of contaminant in autotrophs | g |
| $C_S$ Stock of contaminant in allochthonous autotroph subsidy | g |
| $C_H$ Stock of contaminant in herbivore | g |

|  |  |  |
| --- | --- | --- |
| $E_i$ | Environmental concentration of contaminant from ecosystem $i$,<br>where 1 is local, 0 is donor | $\text{g}\cdot\text{g}^{-1}$ |
| $u_i$ | Environmental uptake of contaminant of trophic level $i$ | $\text{g}\cdot\text{g}^{-1}\cdot\text{time}^{-1}$ |
| $f$ | Assimilation efficiency of the contaminant in the biotic<br>compartment | dimensionless |

Ecosystems, and ecosystem models, have local processes (e.g., feeding interactions among trophic levels) and regional processes that connect ecosystems in space (e.g., fluxes of materials or organisms across ecosystem boundaries) (Massol et al. 2017, Harvey et al. 2021). Here, we focus on spatial ecosystem processes, specifically fluxes of materials connecting ecosystems; for example, via the movement of organisms which transport nutrients to recipient ecosystems (biotic fluxes), or flooding events which lead to the increase or decrease in nutrient inputs in recipient ecosystems (abiotic fluxes). Nutrients and contaminants may, in fact, be transported by similar vectors (though not always) from similar point sources, for example biosolids are used as fertilizers but are also full of contaminants, or leaching and surface runoff of pesticides and fertilizers in agricultural landscapes (Holzem et al. 2014). These vectors can be biotic (e.g., animal migration, ontogenetic habitat shifts) or abiotic (e.g., flooding, wind, water flow). For example, the movement of organisms between ecosystems has been shown to be a critical vector on which contaminants hitch a ride (e.g., Walters et al. 2008, Speir et al. 2014). This includes the yearly spawning run of Chinook salmon transporting nutrients and contaminants from marine environments to upstream freshwater environments as they spawn and die (e.g., Blais et al. 2007). Nutrient inputs, whether abiotic or biotic, also serve as ecological subsidies potentially increasing the primary productivity of the recipient ecosystems (e.g., Polis et al. 1997) with consequent possible effects on contaminant dynamics.

Importantly, there is a potential interaction between these two factors. This has been observed empirically—one of the first studies linking nutrient inputs, or eutrophication, and organic contaminants (DDT, mercury, and PCBs) demonstrated that increasing nutrient inputs to aquatic systems caused an increase in biomass, and thus a dilution of contaminants in the biota, resulting in a lower concentration of pollutants in individual organisms (Olsson and Jensen 1975). This increase in productivity leading to a dilution of contaminants has been termed the bloom-dilution hypothesis (Pickhardt et al. 2002). However, more recent work has shown mixed support for this hypothesis. For example, studies in lake environments have demonstrated that higher nutrient availability promotes growth of phytoplankton and zooplankton, resulting in a decrease in methylmercury concentrations in phytoplankton and zooplankton (Chen and Folt 2000, Pickhardt et al. 2002). In fish populations, however, studies have observed a positive relationship between nutrient loading and mercury concentrations (Driscoll et al. 2012, Chen et al. 2021). In fact, what may be occurring in these cases is that the increase in nutrients is causing indirect effects on higher trophic levels resulting in elevated concentrations of contaminants further up the food web and perhaps other, more complex effects on contaminant distributions in food webs.

It is apparent that there exists empirical evidence to support the hypothesis that ecosystem fluxes (e.g., nutrient inputs) influence contaminant concentrations (e.g., bloom-dilution hypothesis; (Pickhardt et al. 2002). However, general trends are difficult to predict due to the interdependent impacts of the combined dynamics of ecosystems and contaminants. As an initial step in this direction, we derived a novel mathematical framework which integrates ecosystem and contaminant models, and then used this framework to answer three questions: 1) How do spatial ecosystem processes influence contaminant dynamics? 2) How do different types of ecosystem fluxes (or subsidies; Polis et al. 1997), for example within ecosystem fluxes (i.e., the movement of biota from a patch of the same ecosystem type) or across ecosystem fluxes (i.e., the movement of biota from a patch of a different ecosystem type) impact contaminant dynamics? And 3) how do the strengths of different types of ecosystem fluxes impact contaminant dynamics?

### An ecosystem-contaminant coupled model (ECCM)

We couple an ecosystem model with a contaminant model (termed an ecosystem-contaminant coupled model – ECCM hereafter) to examine the impacts of ecosystem fluxes within and across ecosystems on chemical contaminants. In particular, we couple a well-studied ecosystem nutrient model (see Leroux and Loreau 2008) with a novel contaminant model and then use this novel ECCM to investigate the effects of within and across ecosystem fluxes of biotic and/or abiotic materials (*sensu* Massol et al. 2017) on contaminant dynamics. Here, we refer to ecosystem flux as the physical movement of biota or abiotic material from a donor to a recipient ecosystem. This flux can either be within ecosystem or across ecosystems. Specifically, we examine how within ecosystem autotroph and herbivore fluxes (i.e., from a patch of the same ecosystem type; e.g., movement of beetles across two forest patches) and across ecosystem autotroph fluxes (i.e., from a patch of a different ecosystem type; e.g., movement of litter from a riparian forest to a river) influence biomass, contaminant mass, and contaminant concentrations of each trophic level (inorganic nutrients (N), primary producers or autotrophs (A), and primary consumers or herbivores (H)) in the recipient ecosystem.

#### Ecosystem Model

In the simplest case the ecosystem model has two biotic modules: primary producers or autotrophs (A) and primary consumers or herbivores (H), and one abiotic module: inorganic nutrients (N). We use an ecosystem model as it explicitly incorporates the abiotic compartment including abiotic constraints and feedbacks with biotic components—biotic components which are part of food web models. Biomass is then transferred along the food chain in this ecosystem patch through trophic linkages. The recipient ecosystem is open at the basal level through a constant input of inorganic nutrient (*I*, which can be within or across ecosystem types) and a constant loss rate of inorganic nutrient (*l*). This model is then modified to first look at within ecosystem flux by incorporating a constant input of autotrophs or herbivores (*w_A_* or *w_H_*, respectively) whereby immigrating individuals feed and reproduce in the focal patch. And then the model is modified to look at across ecosystem flux by adding a third, donor-controlled biotic module (S) which serves as a subsidy for H but does not directly influence A whereby immigrating individuals do not feed or reproduce in the focal patch but are instead consumed by H (Table 1; Appendix S4: Figure S1). This focus on across ecosystem flux of autotrophs but not herbivores reflects reality since across ecosystem autotroph fluxes (e.g., litter fall) are much more common than across ecosystem herbivore fluxes, making up on average greater than 80% of terrestrial subsidies to aquatic systems (Bartels et al. 2012).

Biotic modules recycle nutrients at rate *d_i_*, where *i* is nutrients (*N*), autotrophs (*A*), or herbivores (*H*), but only a fraction, 1 – δ*_i_*, of the recycled nutrients reaches the soil nutrient pool. Nutrients are recycled via biotic processes such as death, egestion, or excretion, however, there always remains a portion (δ*_i_*) of these nutrients which get lost from the ecosystem. One major route in terrestrial systems is the hydrological loss of dissolved organic molecules during the process of soil humification, leaving the system without remineralization of nutrients and contaminants (Hedin et al. 1995). Likewise, in aquatic systems this includes processes such as sedimentation of detritus (Darchambeau et al. 2005) and hydrological outflow. We make the same simplifying assumption as other ecosystem models that this recycled proportion is instantaneously transformed from organic to inorganic nutrients (i.e., we do not model decomposition or mineralization explicitly, e.g., Loreau 2010, Leroux and Loreau 2012). Finally, similar to other ecosystem models (e.g., Loreau 2010, Leroux and Schmitz 2015) we assume that nutrient uptake can be described using a linear consumption function. In particular, this assumption implies that there will be no upper limits to consumer uptake as would be obtained by incorporating a saturating consumption function. See Appendix S1 for more model details and justifications. The full set of ordinary differential equations representing the ecosystem models presented in Appendix S4: Figure S1 can be found below, where each individual ecosystem model can be recovered by setting the required *w_i_*to 0 (i.e. the base model is recovered by setting *w*_A_ = *w*_S_ = *w*_H_ = 0).

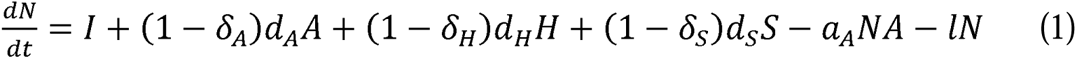

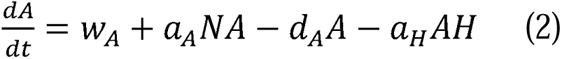

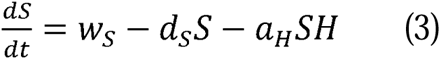

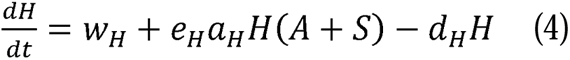

#### Contaminant Model

We model contaminant integration into biota in a similar manner to nutrients with some important differences listed below. The influx of inorganic nutrients at the base of the ecosystem model is adjusted by the environmental concentration of contaminant from the donor ecosystem (thus *I* becomes *I × E_0_*, where *E_0_* is the concentration in the donor ecosystem). We refer to donor ecosystem as the patch from which the ecosystem fluxes originate—for within ecosystem flux this is a donor patch of the same ecosystem type, while for across ecosystem flux this is a donor patch of a different ecosystem type. Similarly, as the ecosystem model becomes modified to look at within– or across-ecosystem flux by incorporating a constant inflow of autotrophs, herbivores, or allochthonous fluxes they are also adjusted by the environmental concentration of contaminant in the donor ecosystem and the environmental uptake rate (*u_i_*) of contaminant for that trophic level (*w_A_ × E_0_× u_A_*, *w_H_ × E_0_× u_H_,* or *w_S_× E_0_× u_S_* respectively; Table 1; Appendix S4: Figure S1). In our model formulation we assume that the influx of inorganic nutrients is from the same ecosystem as the biotic flux (e.g., if it is a within ecosystem biotic flux then it is a within ecosystem influx of inorganic nutrients), however, this assumption could be relaxed in future studies by adding an environmental concentration of contaminant from an alternate donor system (e.g., *E_3_*). Moreover, we assume that our contaminants do not undergo environmental breakdown. This assumption could be relaxed in future studies quite simply by including a contaminant specific loss mechanism for the nutrient compartment.

Unlike a conventional ecosystem nutrient or energy model, biota in the contaminant model are also capable of accumulating contaminants directly from the environment, for example through inhalation, dermal uptake, gill uptake, or foliar uptake (e.g., Devillers 2009). We consider this by an additional term which has an environmental uptake rate of contaminant for trophic level *i* (*u_i_*), which is adjusted by the concentration of the contaminant in the local system (*E_1_*) and the biomass of the trophic level. Not all contaminants can be accumulated directly from their environment (e.g., large polychlorinated biphenyls; McLeod et al. 2015b), and direct environmental uptake can be considered negligible for many contaminant classes for some species (e.g., inhalation for many terrestrial organisms; Smith et al. 2007). In these cases, *u_i_* is equal to zero and this term disappears from the model. This term, *u_i_*, is the equivalent of the parameter *u_i_* within the Ecotracer equations (e.g., Christensen and Walters 2004, McGill et al. 2017) which is described as a parameter representing uptake per biomass per time per unit environmental concentration. Additionally, we assume that contaminants are non-metabolizable.

We assume that biota are also able to accumulate contaminants by ingesting contaminated food, thus the nutrient uptake term from the ecosystem model is adjusted by the concentration of contaminant, *C_i_*, in the given trophic level, *i* (where *i* is autotrophs (*A*), herbivores (*H*)) and an assimilation efficiency of the contaminant in the biotic compartment (*f*). Of course, not all contaminants biomagnify (e.g., some trace metals) – in those cases *f* is equal to zero and this term disappears from the model. Moreover, we assumed that the assimilation efficiency of the contaminant in the biotic compartment is related to chemical properties and not species-specific, however, this assumption could be relaxed in future studies. Our model simulations were carried out for parameter values between 0 and 10 (as per Leroux and Schmitz 2015), inclusive, for all parameters, except δ*_i_*, *e_i_*, and *f* which are proportions constrained between 0 and 1. This is a commonly used wide range of parameter values in ecosystem studies (e.g. Leroux and Schmitz 2015) which allows us to explore general model behaviour while ensuring biological and chemical realism. Consequently, those cases where a contaminant does not biomagnify (i.e., when *f* = 0) are included within our results. This term, *f*, is the equivalent of GC_i_ from the Ecotracer equations (e.g., Christensen and Walters 2004, McGill et al. 2017) which describes the proportion of contaminant assimilated from the ingestion of contaminated food. In this way, terms *f* and *u_i_* incorporate variability in the physicochemical properties of contaminants and could be specified for a representative contaminant (e.g., Appendix S4: Figure S7 & S8). Finally, contaminants can be lost and recycled through the system in the same manner as nutrients. The full set of ordinary differential equations representing the contaminant models presented in Appendix S4: Figure S1 can be found below, where each individual contaminant model can be recovered by setting the required *w_i_*to 0 (i.e. the base model is recovered by setting *w*_A_ = *w*_S_ = *w*_H_ = 0).

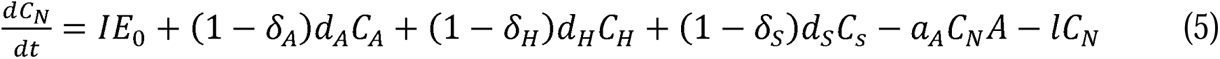

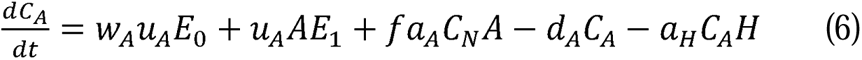

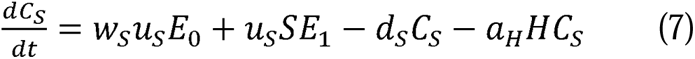

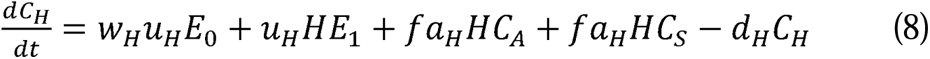

Contaminant concentration is then calculated by dividing the mass of contaminant in a trophic level by the biomass of that trophic level.

### Model Analysis

Using the coupled ecosystem and contaminant model we contrast how within and across ecosystem nutrient fluxes impact contaminant dynamics, specifically compared to a base model where local stock of inorganic nutrients is impacted by nutrient inputs. We considered three broad model types: i) base case (no inflow of biotic fluxes), ii) within ecosystem flux (influx of autotrophs or herbivores), and iii) across ecosystem flux (influx of autotrophs).

#### How do ecosystem processes influence contaminant dynamics in the absence of biotic fluxes?

Using the base case ECCM (i.e., model with no biotic fluxes where *w_A_* = 0 and *w_H_* = 0) we examined whether nutrient input (*I*) and loss (*l*) influenced contaminant concentrations in the biomass of organisms. To do this, we first solve the model for the feasible equilibrium contaminant mass and contaminant concentrations for each trophic level and then we took the partial derivative of each equilibrium value with respect to either nutrient inputs (*I*) or nutrient loss (*l*). Here, equilibrium is when the change in biomass, contaminant mass, or contaminant concentration over time is zero. A positive, negative, and zero partial derivative indicates a positive, negative, or no effect, respectively, of the parameter on equilibrium contaminant mass or concentration. Overall, we aim to investigate general predictions for our ECCM but we also explore some specific contaminant cases. For this question, we investigate how nutrient input and loss impact polychlorinated biphenyls (PCBs) and cyclic methyl siloxanes (CMSs) concentrations – two prominent chemicals in the environment with different chemical uptake chemical properties. PCBs have a higher assimilation efficiency and lower environmental uptake, while CMSs have much lower assimilation efficiency but a higher environmental uptake efficiency (see Appendix S4: Table S1).

#### Do different types of biotic fluxes impact contaminant dynamics differently?

Using the same ECCM, but expanded to include within ecosystem flux and across ecosystem flux, we solved the full model (see Table 1) for equilibria and determined the feasibility conditions for these equilibria (i.e., biomass and contaminant mass must be greater than zero). Then, we randomly selected 10 000 parameters sets in which all parameters were simultaneously varied between 0 and 10 according to a uniform random distribution (as per Leroux and Schmitz 2015) with the exception of δ*_i_*, *e_i_*, and *f* which are constrained between 0 and 1 (see Appendix S1 for additional model justification). We retained only parameters sets that satisfied the feasibility conditions (n = 7951) and used these to calculate numerical equilibria values for each of the three biotic flux scenarios along with the base model scenario.

We compared models by reporting the natural logarithm of the model equilibrium when biotic fluxes are incorporated as compared to the model equilibrium of the base model to determine how biotic fluxes influences biomass, contaminant mass, and contaminant concentrations (see Appendix S4: Figure S2 for conceptual diagram of how to interpret these results). In this way, a value greater than zero implies that biotic fluxes result in a greater biomass, contaminant mass, or contaminant concentration than the base case with no biotic fluxes, while a value less than zero implies that biotic fluxes result in a lower biomass, contaminant mass, or contaminant concentration than the base case.

We then contrasted these results for two specific cases: Case 1 – the recipient ecosystem was more contaminated than the donor ecosystem, i.e., the recipient system is a contaminant hot spot, or Case 2 – the donor ecosystem was more contaminated than the recipient ecosystem, i.e., the donor system is a contaminant hot spot. This was done because results could depend on the direction of the contaminant gradient between recipient and donor ecosystems.

#### How do the strengths of different types of biotic fluxes impact contaminant dynamics?

It has been shown in both theoretical and empirical studies that the impact of biotic fluxes on recipient ecosystems depends on the trophic level that is responsible for the flux (e.g., Leroux and Loreau 2008, Allen and Wesner 2016, Montagano et al. 2019). Thus, it is likely that biotic fluxes by different trophic levels have different relative effects on contaminant mass and contaminant concentrations. Since we explored the consequences of different types of fluxes on equilibria across the same parameter sets, we can do a direct comparison to assess the relative effects of these fluxes on one another. To do this, we used attenuation plots whereby we plotted the model comparison metrics described above against each other for each trophic level (e.g., the natural log of the change in autotroph flux model equilibrium as compared to base model equilibrium vs the natural log of the change in herbivore flux model equilibrium as compared to base model equilibrium). In this way, we determined which subsidy has a stronger impact by examining where the points sit relative to the one-to-one line, with points above the line demonstrating that the subsidy on the y-axis has a stronger impact on equilibria values than those points below the line, and vice versa.

Due to differences in environmental contamination of the donor and recipient ecosystem the subsidy may be moving from a more contaminated ecosystem to a less contaminated ecosystem, or vice versa. It is important to distinguish between these because contaminant concentrations are expected to increase in the recipient ecosystem if the subsidy is coming from a more contaminated system. Thus we contrast four scenarios – Case 1a is when the x and y-axes subsidies are moving from a less contaminated ecosystem to a more contaminated ecosystem, Case 1b is when only the y-axis subsidy is moving from a less contaminated ecosystem to a more contaminated ecosystem, while Case 2a is when the x and y-axes subsidies are moving from a more contaminated ecosystem to a less contaminated ecosystem, and Case 2b is when only the y-axis is moving from a more contaminated ecosystem to a less contaminated ecosystem. Finally, we determined realistic uptake and assimilation efficiency values for different classes of organic chemical contaminants using chemical properties from Walters et al. (2016) and the equations for calculating environmental uptakes (*u_i_*) and assimilation efficiencies (*f*) provided in Arnot and Gobas (2004). We then placed these values on the same plots to determine how biotic fluxes may influence different groups of organic contaminants.

#### Global Sensitivity Analysis

To identify the most influential parameters on our model predictions we performed a global sensitivity analysis (GSA) similar to the one outlined in Bellmore et al. (2014) and Harper et al. (2011). In brief, we used the 10 000 randomly selected parameter sets described before and using the parameter combinations and the predicted contaminant concentrations in each trophic level we applied a random forest algorithm to calculate the residual sum of squared errors for each parameter (Breiman et al. 2011, Bellmore et al. 2014). This is a common method for ranking parameters of ecological models which can then be converted to a relative importance index by normalizing each residual sum of squared errors by the sum total.

## Results

### How do ecosystem processes influence contaminant dynamics in the absence of biotic fluxes?

At equilibrium, biomass for all trophic levels, with the exception of autotrophs (A), depended on both nutrient loss and nutrient inputs. More importantly, when contaminant mass was examined, we observed that contaminant mass and contaminant concentration equilibrium values for all trophic levels depended on both nutrient loss and nutrient input rates (see Appendix S2 for full equilibrium solutions and Appendix S4: Figure S3 for contaminant mass). This was further demonstrated by non-zero partial derivatives for the feasible equilibrium values for each trophic level with respect to both *I* and *l* (see Figure 2).

**Figure 2.**
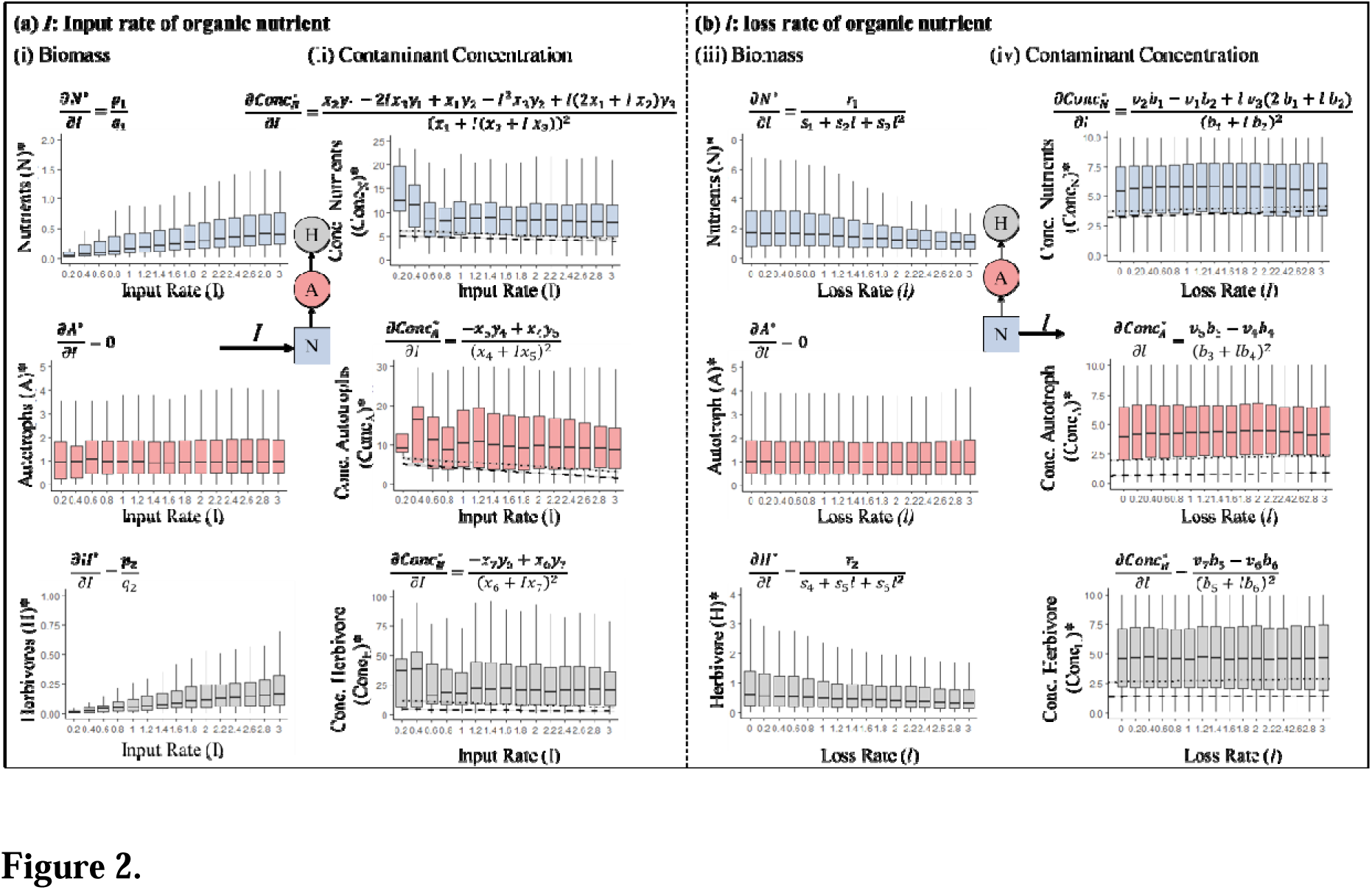
Demonstration of how (a) input rate of organic nutrient (*I*) and (*b*) loss rate of organic nutrient (*l*) influence the (i & iii) equilibrium biomass and (ii & iv) equilibrium contaminant concentration in the absence of biotic fluxes with the range of parameters described in the main text. While these results are dependent on the parameters selected, the simplified partial derivatives below each graph demonstrate that input rate and loss rate influence equilibrium contaminant concentrations irrespective of parameter values. Overlaid on (ii & iv) are two representative chemicals – polychlorinated biphenyls (dotted line) and cyclic methyl siloxanes (dashed line) for the same parameter values, but that have chemical specific values of environmental uptake of contaminant by trophic level *i* (*u_i_*) and assimilation efficiencies of the contaminant in biotic compartment (*f*). For more details on the substitutions for the *p_i_*s, *q_i_*s, *r_i_*s, *s_i_*s, *v_i_*s, *b_i_*s, *x_i_*s, and *y_i_*s see Appendix S3. For full equilibria solutions see Appendix S2 and for demonstration of how input rate of organic nutrient (*I*) and loss rate of organic nutrient (*l*) influence equilibrium contaminant mass see Figure A3.

The non-zero partial derivatives for the feasible equilibrium values for each trophic level with respect to *l* demonstrated that nutrient loss rate impacts contaminant concentrations. Across the range of parameter sets explored, an increase in nutrient loss rate from 0 to 3 resulted in a 50 % decline of the mean contaminant concentration in the nutrient pool, a 10 % decline in the mean contaminant concentration in the autotroph and a 10 % increase in the mean contaminant concentration in the herbivore. The relationship between concentration in the nutrient pool and increasing nutrient loss rate was negative for approximately half of the parameter sets (for example when the recycling rate of the autotroph is very high), while there was a positive relationship between concentration in the autotroph pool and increasing nutrient loss rates for approximately 70 % of the parameter sets (for example when the recycling rate of the autotroph was very low). Despite this wide range of parameter values, our results demonstrate that when chemical parameter values are tailored to specific organic chemical classes (here we used PCBs and CMSs) while preserving the rest of the parameter values we see markedly different relationships for increasing nutrient input rate and increasing nutrient loss rate across trophic level. In particular, we observe a decrease in contaminant concentrations in autotrophs for both organic chemical classes when nutrient input rates are increased, and a greater decrease in concentration in herbivores for PCBs than CMSs. We see similar rates of increase in concentration of both PCBs and CMSs with increasing nutrient loss rate across all trophic levels. This is a rather simplistic view of ecosystems since, as mentioned earlier, ecosystems are not closed systems and it is when ecosystems are linked through time and space that some of the more interesting biomass dynamics occur.

### How do different types of fluxes impact contaminant dynamics?

The incorporation of within or across ecosystem fluxes of any trophic level resulted in differing magnitudes of recipient ecosystem biomass dynamics dependent on the trophic level of the flux moving (Figure 3a). For example, within ecosystem autotroph fluxes resulted in no change in recipient autotroph biomass compared to the base case, while herbivore and across ecosystem autotroph flux both resulted in declines in recipient autotroph biomass compared to the base case, with the incorporation of herbivore flux resulting in a much larger decline in recipient autotroph biomass. We observed similar directional changes in mean contaminant biomass (Figure 3b). For example, all types of fluxes resulted in elevated mean contaminant biomass in both the nutrient pool and the herbivore pool irrespective of the background contaminant concentration of the donor ecosystem. Meanwhile, herbivore and across ecosystem autotroph fluxes both resulted in declines in mean contaminant biomass in the recipient autotroph, again irrespective of the background contaminant concentrations of the donor ecosystem. Importantly, the consistency in the direction of the contaminant biomass results irrespective of the background contaminant concentrations of the donor ecosystem demonstrated that that these results are not merely the result of mixing a highly contaminated ecosystem with a more pristine ecosystem.

**Figure 3.**
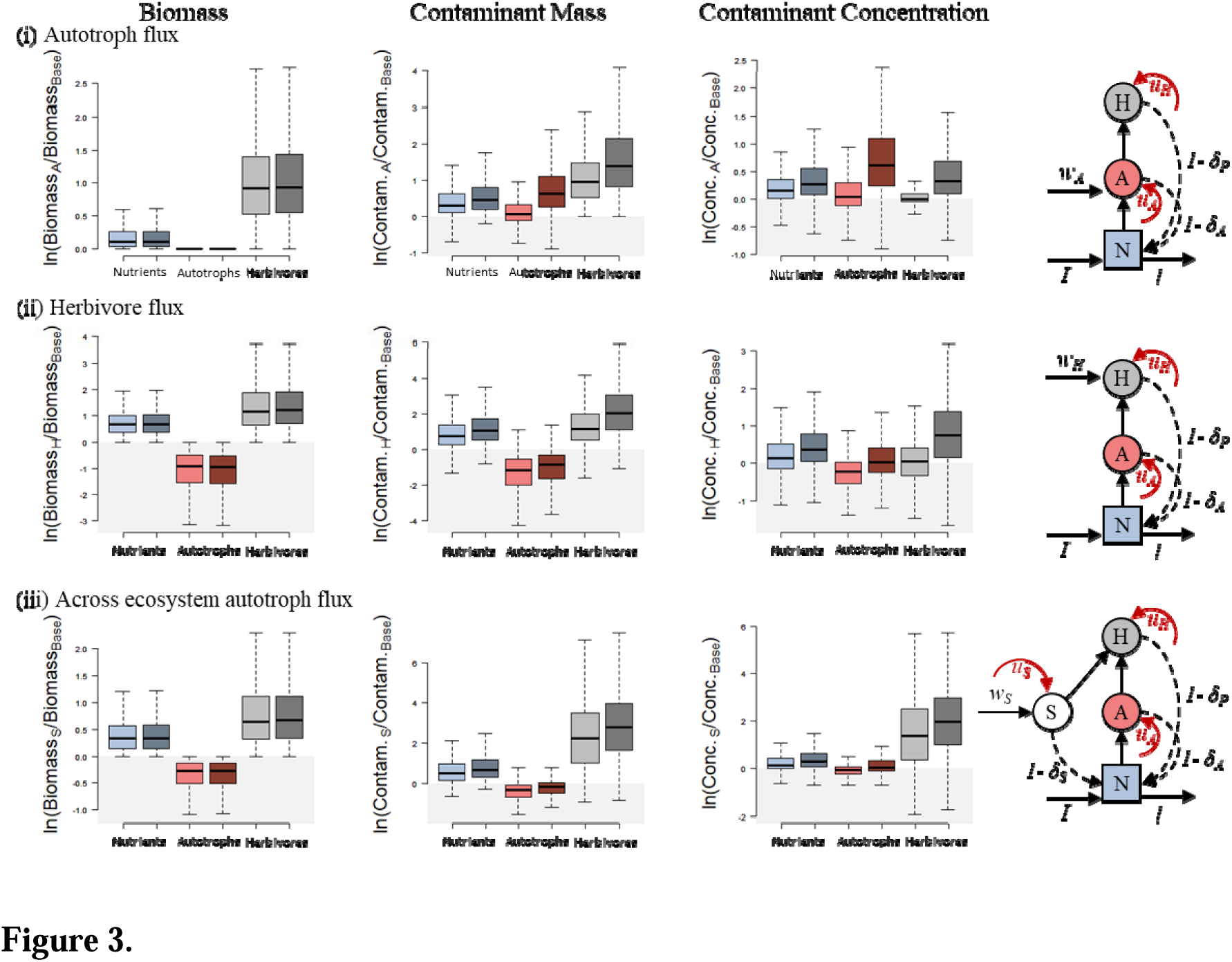
Contaminant concentrations in trophic levels as compared to the no movement base model are influenced by the type of flux (i.e., (i) autotroph fluxes within ecosystems, (ii) herbivore fluxes within ecosystems, and (iii) autotroph fluxes across ecosystems) with the incorporation of fluxes contributing to an increase in the concentration of contaminant in herbivores, irrespective of flux type or contamination level of donor ecosystem. The lighter bars indicate the case in which the recipient ecosystem is more contaminated than the donor ecosystem, and the darker bars indicate that the recipient ecosystem is less contaminated than the donor ecosystem. Columns are grouped as Biomass, Contaminant mass, and Contaminant concentration dynamics and the model diagrams in the last columns are a visual representation of (i), (ii), and (iii). Box plot colours are as follows: blue is inorganic nutrients, red is autotrophs, and grey is herbivores. See Appendix S4: Figure S2 for interpretation of patterns.

The impact of fluxes on contaminant concentrations was then determined by the magnitude with which both the biomass and the contaminant biomass are impacted by the flux. Overall, the incorporation of within or across ecosystems fluxes of any trophic level resulted in elevated concentrations of contaminants in the recipient abiotic nutrient pool irrespective of whether that flux was coming from a more contaminated ecosystem or not (Figure 3c, blue boxes). The variability in nutrient concentration, irrespective of flux type, was dominated by recycling mechanisms, an important ecosystem process, through the dominance of recycling rate of the herbivore (*d_H_*) and proportion of material lost by the herbivore (δ*_H_*) and several contaminant processes including how contaminated the donor system was (*E_0_*) and how well the organism could assimilate the contaminant (*f*) (Figure 4). Fluxes impacted the contaminant concentrations in the autotrophs more predictably – when the flux was from a donor system which was more pristine than the recipient system, the flux resulted in a decrease in recipient autotroph concentration with the opposite being true when the flux was from a more contaminated ecosystem (Figure 3c red boxes). This was apparent from the global sensitivity analysis as well, where the contaminant parameters were the dominant drivers in recipient autotroph contaminant concentrations (Figure 4). Finally, the incorporation of biotic fluxes resulted in increased contaminant concentrations in recipient herbivores for most parameter sets in most scenarios, irrespective of the contamination level of the donor ecosystem (Figure 3c). The parameter driving this elevated concentration in herbivores was flux dependent. Consumption processes, i.e., organism assimilation efficiency (*e_H_*) was the driving parameter when there was autotroph flux (driving over 20% of herbivore contaminant concentrations). On the other hand, when there was herbivore flux, recycling processes, i.e., the proportion of material lost by the herbivore (δ_H_) was the most important parameter (driving close to 20% of the variation in herbivore contaminant concentrations; see Figure 4).

**Figure 4.**
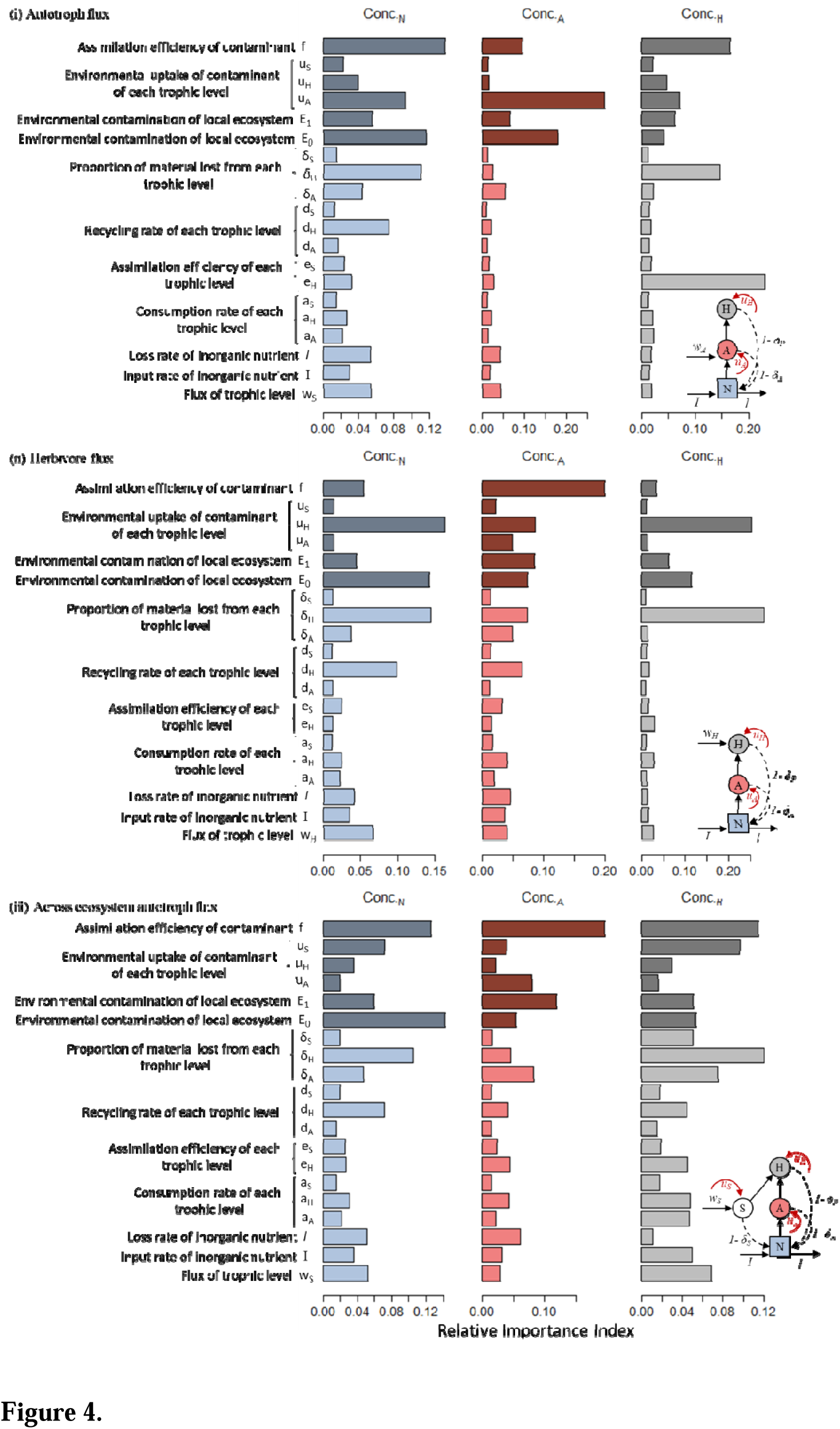
Relative importance index from the global sensitivity analysis examining the importance of parameters on contaminant concentrations in each trophic level when there is (i) autotroph flux, (ii) herbivore flux, and (iii) across ecosystem autotroph flux. This figure demonstrates the importance of consumptive (e.g., *e_H_*) and recycling (e.g., δ*_H_*) processes for nutrient and herbivore contaminant concentrations and contaminant parameters for autotroph contaminant concentrations. In each panel the colours represent the parameter contributions to nutrient concentrations (blue), autotroph concentrations (red), and herbivore concentrations (grey). The darker bars indicate parameters from the contaminant model, while the lighter bars highlight ecosystem parameters.

### How do the strengths of different types of biotic fluxes impact contaminant dynamics?

Autotroph fluxes within ecosystems had a stronger impact on recipient autotroph contaminant concentrations than autotroph fluxes across ecosystems (Figure 5 ii, middle), however, both had a stronger impact than herbivore fluxes irrespective of environmental contamination (Figure 5 ii). These results were similar to the observations for biomass and contaminant mass, albeit weaker.

**Figure 5.**
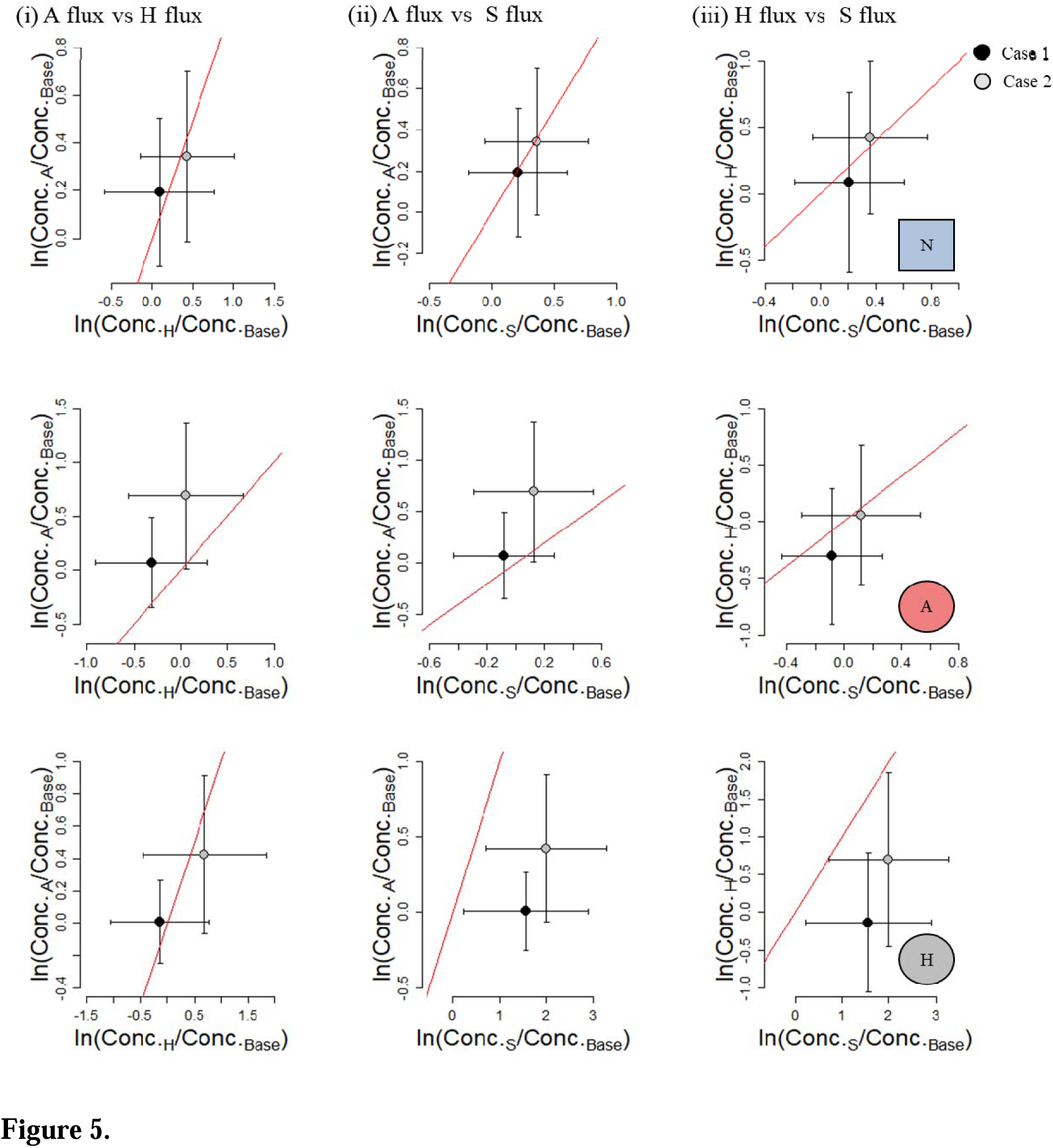
Attenuation plots comparing the influence of (i) within ecosystem autotroph fluxes on contaminant concentration to within ecosystem herbivore fluxes, (ii) within ecosystem autotroph fluxes to across ecosystem autotroph fluxes, and (iii) within ecosystem herbivore fluxes to across ecosystem autotroph fluxes. Plots demonstrate that the type of flux has little impact on nutrient contaminant concentration (top row), while within and across ecosystem autotroph flux had a stronger impact on contaminant concentration in autotrophs than herbivore subsidies (middle row), and that across ecosystem autotroph flux had a stronger impact on contaminant concentration in herbivores than other types of flux. Case 1 is when the recipient ecosystem is more contaminated than the donor ecosystem, and Case 2 is when the recipient ecosystem is less contaminated than the donor ecosystem. The red line indicates a one-to-one relationship and everything above and to the left of the red line means the y-axis has a stronger impact than the x-axis, while everything below and to the right of the red line means the x-axis has a stronger impact than the y-axis (see Appendix S4: Figure S4 for biomass and Appendix S4: Figure S7 for contaminant mass). The dots are the mean value from simulations with standard deviation confidence bars. See Appendix S4: Figure S8 for similar figure with organic chemical classes plotted.

The effects of the influx of herbivores were never overwhelmingly stronger than the influx of autotrophs from within ecosystems for recipient herbivore concentrations, however between ecosystem autotroph fluxes had a stronger impact on recipient herbivore concentrations than within ecosystem herbivore fluxes. Moreover, autotroph fluxes across ecosystems had a stronger impact on *in situ* herbivore concentrations than within ecosystem autotroph fluxes, irrespective of environmental contamination (Figure 5 ii, bottom).

Finally, when thinking about specific organic chemical classes, we see that values for the chemical specific properties (i.e., *f* and *u_i_*) of all the organic chemicals have ranges within the parameter ranges chosen for the general simulations. Moreover, for many organic chemical classes these ranges overlap (e.g., assimilation efficiency (*f*) ranges from 0.004 to 0.404 for Brominated Flame Retardants, and from 0.002 to 0.479 for Polychlorinated Biphenyls (PCBs); Appendix S4: Table S1). Cyclic Methyl Siloxanes (CMS), however, have much lower assimilation efficiency ranges (0.003–0.049) but a higher environmental uptake efficiency (maximum is 0.542). Thus, CMS behaves slightly differently, a deviation which was most pronounced in the herbivore plots (Appendix S4: Figure S8), where herbivore movement was more influential on CMS concentrations than between ecosystem autotroph movement which was in direct contrast with all other classes of organic chemicals examined here.

## Discussion

By coupling an ecosystem model with a contaminant model, we mathematically demonstrate the relationship between contaminant concentrations and spatial ecosystem processes from nutrient input and loss rates to biotic fluxes of different trophic levels. In particular, we demonstrate two key results (1) the role of ecosystem processes, for example increasing nutrient loss rate leads to increasing contaminant concentrations across trophic levels, and the importance of ecosystem properties such as recycling on contaminant concentrations, (2) the role of movement, particularly of lower trophic levels, on increasing herbivore contaminant concentrations. Moreover, these results are largely conserved across a broad range of chemical classes simulated (e.g. Alkylphenols, CMSs, PCBs). We finish by demonstrating the further application of these results and the ECCM approach for (3) remediation efforts and (4) exploring more complex ecosystem and contaminant dynamics.

### Ecosystem processes affect contaminant concentrations

It has been demonstrated both empirically and theoretically that biomass of a trophic level depends on both nutrient input rates and loss rates (e.g., Oksanen et al. 1981, Leibold 1996) which, in turn, should alter the concentration of contaminants in a trophic level. However, empirical support for this hypothesis (the bloom-dilution hypothesis; *sensu* Pickhardt et al. 2002) is mixed, with studies typically finding a large amount of unexplained variation in the relationship between nutrient inputs and concentrations of contaminant in biota (Pickhardt et al. 2002). Environmental systems, however, are complex – for example, even mesocosm studies, which offer more control over experimental design and potential confounding factors, can have surprising results. In a synthesis of recent mesocosm studies on how elevated nitrogen and phosphorus inputs impacted methylmercury fluxes from aquatic to terrestrial habitats, (Chumchal and Drenner 2020) observed that nutrient addition led to an increase in primary producer biomass and a concurrent increase in insect-mediated methylmercury flux from small midges, but that methylmercury concentrations in dragonflies and large midges were largely unaffected. They argued that the impact of nutrient addition on contaminant concentrations may have been masked by unintuitive changes in biomass at different trophic levels due to ecosystem effects resulting from nutrient additions (e.g., trophic cascades; Chumchal and Drenner 2020).

Our model helps to reconcile this empirical uncertainty by highlighting the importance of ecosystem processes, not just nutrients, on contaminant concentrations in biota. For example, recycling of nutrients is an important process governing concentration of contaminants in herbivores, particularly when there is an across ecosystem autotroph flux, or a within ecosystem herbivore flux (Figure 4). These results suggest that measuring ecosystem properties such as recycling (i.e., senescence, excretion, decomposition, mineralization) could be critical for understanding the context-dependent drivers of variation in the bloom-dilution hypothesis.

Moreover, our model demonstrates how unintuitive impacts of trophic fluxes on contaminant concentrations emerge. For example, we observe that herbivore fluxes within an ecosystem contribute very little to contaminant concentrations in the abiotic pool as compared to within and across ecosystem autotroph flux (Figure 5 i & iii, top). Yet, biomass and contaminant mass demonstrated the opposite results – that is, herbivores fluxes had a stronger impact on both biomass and contaminant mass than either within or across ecosystem autotroph fluxes irrespective of background contamination (Appendix S4: Figure S7). Instead, the dynamics in both biomass and contaminant biomass negate each other resulting in a greatly dampened impact of herbivore fluxes on contaminant concentrations as compared to autotroph fluxes within or across ecosystems. We explore the coupling of a contaminant-ecosystem model within a very simple ecosystem; nevertheless, complex dynamics still emerge. Future work, however, is critical for exploring how more complex ecosystems will alter our general predictions including the incorporation of vertical (e.g., adding a predator) or horizontal (e.g., adding an additional plant compartment) diversity. Incorporating species traits (e.g., body size) into future models may capture key aspects of the ecology while maintaining a relatively simple model structure (Schmitz and Leroux 2020).

### Subsidy dynamics impact contaminant concentrations

Subsidies have a strong impact on contaminant concentrations, irrespective of what is moving (Polis et al. 1997). For example, Walters et al. (2008) estimated that the emergence of adult insects in a 25 km riparian reach exports the same mass of polychlorinated biphenyls as the amount deposited by 50 000 Chinook salmon when they spawn in the Pacific Northwest of the United States. What our model highlights, however, is that ecosystem fluxes can impact more than just the movement of contaminants as they hitch rides on biotic vectors. In particular, the evidence that incorporating within ecosystem autotroph fluxes contributes to a higher concentration of chemical contaminant in a consumer trophic level (herbivores) highlights the importance of movement, especially of lower trophic levels, on recipient ecosystems. We modeled subsidies as continuous fluxes, however, empirical evidence shows that many subsidies (e.g. emergent insects, spawning salmonids) are temporally variable (Bartels et al. 2012, McCary et al. 2021). Future models could incorporate temporal dynamics in subsidy flux (see Takimoto et al. 2009, Leroux and Loreau 2012, McCary et al. 2021) to investigate how such dynamics impact recipient ecosystem contaminant concentrations through time. Moreover, while our results demonstrate that, on average, cross-ecosystem autotroph fluxes led to an increase in herbivore contaminant concentrations for a range of contaminants from large, hydrophobic compounds which biomagnify, to more water-soluble contaminants for which dietary exposure is not a dominant uptake route, the importance of these aquatic-terrestrial linkages is contaminant dependent (e.g., Liu et al. 2021). For example, trace metals can be lost during the metamorphosis by amphibians (e.g., Snodgrass et al. 2003) and aquatic insects (e.g., Kraus et al. 2021b) resulting in minimal cross-ecosystem transfer of trace metal contaminants. In particular, the large variation in our results (e.g., Figure 3 iii) demonstrates the sensitivity of these results to chemical-specific parameters including both assimilation efficiency of the contaminant and the environmental uptake rates (see Figure 4), highlighting how our ECCM could be used to explore these empirical results further with a more tailored analyses of those contaminants.

### Perspectives on future models and applications in remediation

Teasing out the relationships between nutrient inputs and contaminant concentrations is particularly critical as remediation strategies typically focus on decreasing background contaminant concentrations (e.g., Gobas and Arnot 2010). If changes to nutrient input and loss rates can influence contaminant concentrations in biota, however, then the increase in nutrient stresses due to anthropogenic change (e.g., cultural eutrophication) may also impact contaminant concentrations. In particular, at least in a system without biotic fluxes, our results appear to support the bloom-dilution hypothesis, whereby increasing nutrient inputs causes an increase in biomass and, in the absence of a concurrent increase in contaminant inputs, a decrease in contaminant concentrations. For example, despite no change in autotroph (A) biomass with increasing nutrient input, a decrease in concentration of contaminant in A was observed with increasing nutrient input rates (Figure 2). One remediation technique is phytoremediation and extraction where nutrients are added to contaminated soil, often mine tailings, to promote plant growth. The plants then accumulate the contaminant in the harvestable parts of the plant and this is disposed of (see review by Wang et al. 2017). Our results demonstrate that ecosystem-contaminant coupled models, as the one we derive, could be used to optimize nutrient addition to maximize growth and minimize the dilution effect, to ensure the most cost-effective harvest of contaminated plant matter. This is a bottom-up approach to remediation, however, changes in the biomass of different trophic levels can result from consumer processes at the top of ecosystems. For example, it has been shown that grazing pressure of snails on biofilm communities led to elevated concentrations of atrazine in biofilm compared to a control group without snails (Muñoz et al. 2001). This is an area where an ECCM is particularly useful – by coupling an ecosystem model with a contaminant model we can explore the unexpected direct and indirect impacts that adding an alternate consumer (increasing horizontal complexity) or adding a predator (increasing vertical complexity) might have on coupled biotic and abiotic components of ecosystems.

The framework presented here can be tailored beyond just chemical-specific parameters to one which incorporates more complex contaminant dynamics. Indeed, the production of synthetic chemicals globally has grown exponentially in recent decades (Rockström et al. 2009, Bernhardt et al. 2017) and these chemicals are often designed with specific purposes in mind such as acute, deleterious effects on insects deemed pest species by agriculture. Despite the obvious direct ecological implications of this proliferation in chemical production, from extirpating key species in communities (e.g., Köhler and Triebskorn 2013) to altering population growth rates of top predators (e.g., DDT and peregrine falcons; Hellou et al. 2013), ecological research examining the synergistic effects of contaminants on ecosystems has remained infrequent (Bernhardt et al. 2017). In Appendix S6 we demonstrate how modifying mortality rate of a given trophic compartment to being a function of contaminant concentration can impact equilibrium biomass and contaminant concentrations. Recent studies have examined chemical flux by aquatic-terrestrial linkages, or mass of chemical transported by adult emerging insects, on gradients of donor ecosystem contamination (e.g., Kraus et al. 2021a). In particular, Kraus et al. 2021a observed that higher mortality rates in highly contaminated donor ecosystems leads to a lower emergence of aquatic insects and thus a lower contaminant mass being transported out of the system. Our framework is an important next step in considering how ecosystems may affect contaminant concentrations – a critical avenue for remediation and human health concerns in the face of a changing environment – and on how contaminants may influence ecosystem processes.

## Conclusions

In this work we advanced a relatively simple framework to demonstrate how spatial ecosystem processes (e.g., inorganic nutrient and biotic fluxes) can have unintuitive impacts on contaminant concentrations in an ecosystem with a simple food chain. We demonstrated that the complex interplay between contaminant concentrations and ecosystem stressors is undeniable. In particular, we demonstrated how fluxes can impact contaminant dynamics and contrasted the relative effects of biotic fluxes on contaminants, highlighting the importance of nutrient recycling stressing the need for an ecosystem coupled contaminant model. This is particularly relevant in aquatic ecosystems which are likely to receive chemical contaminants and allochthonous resources through the downhill movement of water, sediment, detritus and associated contaminants (Allan 2004). However, there is a need in both aquatic and terrestrial systems for a) understanding both the impact of ecosystem fluxes on contaminant bioaccumulation, for example for more effective remediation strategies, and b) understanding how contaminants can influence ecosystem structure and function, for example for anticipating ecosystem change. Unfortunately, the current approach to understanding contaminant impacts through tests of single compounds for acute effects on single organisms does not provide insights into the indirect effects contaminants may have on ecosystems (see further discussion in Bernhardt et al. 2017) through chronic effects such as skewed sex ratios, or reduced prey abundance leading to trophic cascades (e.g., Halstead et al. 2014, Rogers et al. 2016). Thus, an ecosystem approach to studying the effects of contaminant dynamics is critical going forward. One major driver of anthropogenic change is habitat destruction and the consequential fragmentation of ecosystems decreasing the movement ability of organisms (Tucker et al. 2018), including changes to aquatic habitat connectivity such as damming of rivers (Grill et al. 2015). Incorporating movement of biota between communities in ecotoxicology models is still in its infancy (see McLeod et al. 2015a, Li et al. 2019), however, these results demonstrate the importance of ecosystem linkages for understanding local contaminant dynamics.

## Acknowledgements

This research was funded by an NSERC Discovery Grant to SJL, an NSERC PGS-D scholarship to AMM, a FAPESP Grant (São Paulo Research Foundation: 2015/18790-3) to LS, and an US-NSF Grant (2025118) to MAL.

## References

1. Allan, J. D. 2004. Landscapes and riverscapes: the influence of land use on stream ecosystems. Annu. Rev. Ecol. Evol. Syst. 35:257–284.

2. Allen, D. C., and J. S. Wesner. 2016. Synthesis: comparing effects of resource and consumer fluxes into recipient food webs using metalJanalysis. Ecology 97:594–604.

3. Arnot, J. A., and F. A. P. C. Gobas. 2004. A food web bioaccumulation model for organic chemicals in aquatic ecosystems. Environmental Toxicology and Chemistry 23:2343– 2355.

4. Bartels, P., J. Cucherousset, K. Steger, P. Eklöv, L. J. Tranvik, and H. Hillebrand. 2012. Reciprocal subsidies between freshwater and terrestrial ecosystems structure consumer resource dynamics. Ecology 93:1173–1182.

5. Bellmore, J. R., A. K. Fremier, F. Mejia, and M. Newsom. 2014. The response of stream periphyton to P acific salmon: using a model to understand the role of environmental context. Freshwater Biology 59:1437–1451.

6. Bernhardt, E. S., E. J. Rosi, and M. O. Gessner. 2017. Synthetic chemicals as agents of global change. Frontiers in Ecology and the Environment 15:84–90.

7. Blais, J. M., R. W. Macdonald, D. Mackay, E. Webster, C. Harvey, and J. P. Smol. 2007. Biologically mediated transport of contaminants to aquatic systems. Environmental Science & Technology 41:1075–1084.

8. Breiman, L., A. Cutler, A. Liaw, and M. Wiener. 2011. Package ‘randomforest.’ Software available at: http://stat-www.berkeley.edu/users/breiman/RandomForests.

9. Burton Jr, G. A., R. Di Giulio, D. Costello, and J. R. Rohr. 2017. Slipping through the cracks: why is the US Environmental Protection Agency not funding extramural research on chemicals in our environment? Environmental Science & Technology 51:755–756.

10. Chen, C. Y., K. L. Buckman, A. Shaw, A. Curtis, M. Taylor, M. Montesdeoca, and C. Driscoll. 2021. The influence of nutrient loading on methylmercury availability in long island estuaries. Environmental Pollution 268:115510.

11. Chen, C. Y., and C. L. Folt. 2000. Bioaccumulation and diminution of arsenic and lead in a freshwater food web. Environmental Science & Technology 34:3878–3884.

12. Christensen, V., and C. J. Walters. 2004. Ecopath with Ecosim: methods, capabilities and limitations. Ecological modelling 172:109–139.

13. Chumchal, M. M., and R. W. Drenner. 2020. Ecological Factors Controlling Insect-Mediated Methyl Mercury Flux from Aquatic to Terrestrial Ecosystems: Lessons Learned from Mesocosm and Pond Experiments. Pages 17–33 Contaminants and Ecological Subsidies. Springer.

14. Darchambeau, F., I. Thys, B. Leporcq, L. Hoffmann, and J. Descy. 2005. Influence of zooplankton stoichiometry on nutrient sedimentation in a lake system. Limnology and Oceanography 50:905–913.

15. Devillers, J. 2009. Ecotoxicology modeling. Springer.

16. Driscoll, C. T., C. Y. Chen, C. R. Hammerschmidt, R. P. Mason, C. C. Gilmour, E. M. Sunderland, B. K. Greenfield, K. L. Buckman, and C. H. Lamborg. 2012. Nutrient supply and mercury dynamics in marine ecosystems: A conceptual model. Environmental Research 119:118–131.

17. Gobas, F. A., and J. A. Arnot. 2010. Food web bioaccumulation model for polychlorinated biphenyls in San Francisco Bay, California, USA. Environmental Toxicology and Chemistry 29:1385–1395.

18. Gounand, I., E. Harvey, C. J. Little, and F. Altermatt. 2018. Meta-ecosystems 2.0: rooting the theory into the field. Trends in Ecology & Evolution 33:36–46.

19. Gravel, D., F. Guichard, M. Loreau, and N. Mouquet. 2010. Source and sink dynamics in metalJecosystems. Ecology 91:2172–2184.

20. Grill, G., B. Lehner, A. E. Lumsdon, G. K. MacDonald, C. Zarfl, and C. R. Liermann. 2015. An index-based framework for assessing patterns and trends in river fragmentation and flow regulation by global dams at multiple scales. Environmental Research Letters 10:015001.

21. Halstead, N. T., T. A. McMahon, S. A. Johnson, T. R. Raffel, J. M. Romansic, P. W. Crumrine, and J. R. Rohr. 2014. Community ecology theory predicts the effects of agrochemical mixtures on aquatic biodiversity and ecosystem properties. Ecology Letters 17:932–941.

22. Hanski, I. 1998. Metapopulation dynamics. Nature 396:41–49.

23. Harper, E. B., J. C. Stella, and A. K. Fremier. 2011. Global sensitivity analysis for complex ecological models: a case study of riparian cottonwood population dynamics. Ecological Applications 21:1225–1240.

24. Harvey, E., I. Gounand, E. A. Fronhofer, and F. Altermatt. 2020. Metaecosystem dynamics drive community composition in experimental, multilJlayered spatial networks. Oikos 129:402–412.

25. Harvey, E., I. Gounand, C. L. Ward, and F. Altermatt. 2017. Bridging ecology and conservation: from ecological networks to ecosystem function. Journal of applied ecology 54:371–379.

26. Harvey, E., J. N. Marleau, I. Gounand, S. J. Leroux, C. Firkowski, F. Altermatt, F. G. Blanchet, K. Cazelles, C. Chu, C. d’Aloia, L. Donelle, D. Gravel, F. Guichard, K. S. McCann, J. Ruppert, C. L. Ward, and M.-J. Fortin. 2021. A general meta-ecosystem model to predict ecosystem function at landscape extents. HAL open science hal-03407501.

27. Hedin, L. O., J. J. Armesto, and A. H. Johnson. 1995. Patterns of nutrient loss from unpolluted, oldlJgrowth temperate forests: Evaluation of biogeochemical theory. Ecology 76:493– 509.

28. Hellou, J., M. Lebeuf, and M. Rudi. 2013. Review on DDT and metabolites in birds and mammals of aquatic ecosystems. Environmental Reviews 21:53–69.

29. Holzem, R., H. Stapleton, and C. Gunsch. 2014. Determining the ecological impacts of organic contaminants in biosolids using a high-throughput colorimetric denitrification assay: a case study with antimicrobial agents. Environmental Science & Technology 48:1646– 1655.

30. Köhler, H.-R., and R. Triebskorn. 2013. Wildlife ecotoxicology of pesticides: can we track effects to the population level and beyond? Science 341:759–765.

31. Kraus, J. M., K. M. Kuivila, M. L. Hladik, N. Shook, D. M. Mushet, K. Dowdy, and R. Harrington. 2021a. CrosslJecosystem fluxes of pesticides from prairie wetlands mediated by aquatic insect emergence: implications for terrestrial insectivores. Environmental Toxicology and Chemistry 40:2282–2296.

32. Kraus, J. M., R. B. Wanty, T. S. Schmidt, D. M. Walters, and R. E. Wolf. 2021b. Variation in metal concentrations across a large contamination gradient is reflected in stream but not linked riparian food webs. Science of The Total Environment 769:144714.

33. Kraus, J. M., J. Wesner, and D. M. Walters. 2020. Synthesis: a framework for predicting the dark side of ecological subsidies. Pages 343–372 Contaminants and Ecological Subsidies. Springer.

34. Larsen, S., J. D. Muehlbauer, and E. Marti. 2016. Resource subsidies between stream and terrestrial ecosystems under global change. Global Change Biology 22:2489–2504.

35. Leibold, M. A. 1996. A graphical model of keystone predators in food webs: trophic regulation of abundance, incidence, and diversity patterns in communities. The American Naturalist 147:784–812.

36. Leroux, S. J., and M. Loreau. 2008. Subsidy hypothesis and strength of trophic cascades across ecosystems. Ecology letters 11:1147–1156.

37. Leroux, S. J., and M. Loreau. 2012. Dynamics of reciprocal pulsed subsidies in local and meta-ecosystems. Ecosystems 15:48–59.

38. Leroux, S. J., and O. J. Schmitz. 2015. PredatorlJdriven elemental cycling: The impact of predation and risk effects on ecosystem stoichiometry. Ecology and Evolution 5:4976– 4988.

39. Li, J., A. M. Mcleod, S. P. Bhavsar, J. Bohr, A. GrgicaklJMannion, and K. Drouillard. 2019. Use of a Food Web Bioaccumulation Model to Uncover Spatially Integrated Polychlorinated Biphenyl Exposures in Detroit River Sport Fish. Environmental Toxicology and Chemistry 38:2771–2784.

40. Liu, Y.-E., X.-J. Luo, Y. Liu, Y.-H. Zeng, and B.-X. Mai. 2021. Bioaccumulation of legacy and emerging organophosphorus flame retardants and plasticizers in insects during metamorphosis. Journal of Hazardous Materials 406:124688.

41. Loreau, M. 2010. From populations to ecosystems. Princeton University Press.

42. Loreau, M., N. Mouquet, and R. D. Holt. 2003. MetalJecosystems: a theoretical framework for a spatial ecosystem ecology. Ecology Letters 6:673–679.

43. Malaj, E., C. Peter, M. Grote, R. Kühne, C. P. Mondy, P. Usseglio-Polatera, W. Brack, and R. B. Schäfer. 2014. Organic chemicals jeopardize the health of freshwater ecosystems on the continental scale. Proceedings of the National Academy of Sciences 111:9549–9554.

44. Marleau, J. N., T. Peller, F. Guichard, and A. Gonzalez. 2020. Converting Ecological Currencies: Energy, Material, and Information Flows. Trends in Ecology & Evolution 35:1068–1077.

45. Massol, F., F. Altermatt, I. Gounand, D. Gravel, M. A. Leibold, and N. Mouquet. 2017. How lifelJhistory traits affect ecosystem properties: effects of dispersal in metalJecosystems. Oikos 126:532–546.

46. Massol, F., D. Gravel, N. Mouquet, M. W. Cadotte, T. Fukami, and M. A. Leibold. 2011. Linking community and ecosystem dynamics through spatial ecology. Ecology Letters 14:313–323.

47. McCann, K. S., K. Cazelles, A. S. MacDougall, G. F. Fussmann, C. Bieg, M. Cristescu, J. M. Fryxell, G. Gellner, B. Lapointe, and A. Gonzalez. 2021. Landscape modification and nutrientlJdriven instability at a distance. Ecology Letters 24:398–414.

48. McCary, M. A., J. S. Phillips, T. Ramiadantsoa, L. A. Nell, A. R. McCormick, and J. C. Botsch. 2021. Transient toplJdown and bottomlJup effects of resources pulsed to multiple trophic levels. Ecology 102:e03197.

49. McGill, L. M., B. S. Gerig, D. T. Chaloner, and G. A. Lamberti. 2017. An ecosystem model for evaluating the effects of introduced Pacific salmon on contaminant burdens of stream-resident fish. Ecological Modelling 355:39–48.

50. McLeod, A. M., J. A. Arnot, K. Borgå, H. Selck, D. R. Kashian, A. Krause, G. Paterson, G. D. Haffner, and K. G. Drouillard. 2015a. Quantifying uncertainty in the trophic magnification factor related to spatial movements of organisms in a food web: Uncertainty and the Effect of Spatial Movements on TMFs. Integrated Environmental Assessment and Management 11:306–318.

51. McLeod, A. M., G. Paterson, K. G. Drouillard, and G. D. Haffner. 2015b. PCB Food Web Dynamics Quantify Nutrient and Energy Flow in Aquatic Ecosystems. Environmental Science & Technology 49:12832–12839.

52. Menge, B. A., T. C. Gouhier, S. D. Hacker, F. Chan, and K. J. Nielsen. 2015. Are metalJecosystems organized hierarchically? A model and test in rocky intertidal habitats. Ecological Monographs 85:213–233.

53. Montagano, L., S. J. Leroux, M. Giroux, and N. Lecomte. 2019. The strength of ecological subsidies across ecosystems: a latitudinal gradient of direct and indirect impacts on food webs. Ecology letters 22:265–274.

54. Muehlbauer, J. D., S. Larsen, M. Jonsson, and E. J. Emilson. 2020. Variables affecting resource subsidies from streams and rivers to land and their susceptibility to global change stressors. Pages 129–155 Contaminants and Ecological Subsidies. Springer.

55. Muñoz, I., M. Real, H. Guasch, E. Navarro, and S. Sabater. 2001. Effects of atrazine on periphyton under grazing pressure. Aquatic Toxicology 55:239–249.

56. Nelson, G. 2005. Drivers of ecosystem change: summary chapter. Ecosystems and human well-being: current state and trends. Millennium Ecosystem Assessment.

57. Oksanen, L., S. D. Fretwell, J. Arruda, and P. Niemela. 1981. Exploitation ecosystems in gradients of primary productivity. The American Naturalist 118:240–261.

58. Olsson, M., and S. Jensen. 1975. Pike as the test organism for mercury, DDT and PCB pollution. A study of the contamination in the Stockholm archipelago. Rep Inst Freshwater Res Drottningholm Fish Board Swed.

59. Pickhardt, P. C., C. L. Folt, C. Y. Chen, B. Klaue, and J. D. Blum. 2002. Algal blooms reduce the uptake of toxic methylmercury in freshwater food webs. Proceedings of the National Academy of Sciences 99:4419–4423.

60. Polis, G. A., W. B. Anderson, and R. D. Holt. 1997. Toward an integration of landscape and food web ecology: the dynamics of spatially subsidized food webs. Annual Review of Ecology and Systematics 28:289–316.

61. Rockström, J., W. Steffen, K. Noone, Å. Persson, F. S. Chapin, E. F. Lambin, T. M. Lenton, M. Scheffer, C. Folke, and H. J. Schellnhuber. 2009. A safe operating space for humanity. Nature 461:472–475.

62. Rogers, H. A., T. S. Schmidt, B. L. Dabney, M. L. Hladik, B. J. Mahler, and P. C. Van Metre. 2016. Bifenthrin causes trophic cascade and altered insect emergence in mesocosms: implications for small streams. Environmental Science & Technology 50:11974–11983.

63. Schiesari, L., M. A. Leibold, and G. A. Burton Jr. 2018. Metacommunities, metaecosystems and the environmental fate of chemical contaminants. Journal of Applied Ecology 55:1553– 1563.

64. Schmitz, O. J., and S. J. Leroux. 2020. Food webs and ecosystems: linking species interactions to the carbon cycle. Annual Review of Ecology, Evolution, and Systematics 51:271–295.

65. Smith, P. N., G. P. Cobb, C. Godard-Codding, D. Hoff, S. T. McMurry, T. R. Rainwater, and K. D. Reynolds. 2007. Contaminant exposure in terrestrial vertebrates. Environmental Pollution 150:41–64.

66. Snodgrass, J. W., W. A. Hopkins, and J. H. Roe. 2003. Relationships among developmental stage, metamorphic timing, and concentrations of elements in bullfrogs (Rana catesbeiana). Environmental Toxicology and Chemistry 22:1597–1604.

67. Speir, S. L., M. M. Chumchal, R. W. Drenner, W. G. Cocke, M. E. Lewis, and H. J. Whitt. 2014. Methyl mercury and stable isotopes of nitrogen reveal that a terrestrial spider has a diet of emergent aquatic insects. Environmental toxicology and chemistry 33:2506–2509.

68. Takimoto, G., T. Iwata, and M. Murakami. 2009. Timescale hierarchy determines the indirect effects of fluctuating subsidy inputs on in situ resources. The American Naturalist 173:200–211.

69. Tucker, M. A., K. Böhning-Gaese, W. F. Fagan, J. M. Fryxell, B. Van Moorter, S. C. Alberts, A. H. Ali, A. M. Allen, N. Attias, and T. Avgar. 2018. Moving in the Anthropocene: Global reductions in terrestrial mammalian movements. Science 359:466–469.

70. Walters, D., T. Jardine, B. S. Cade, K. Kidd, D. Muir, and P. Leipzig-Scott. 2016. Trophic magnification of organic chemicals: A global synthesis. Environmental Science & Technology 50:4650–4658.

71. Walters, D. M., K. M. Fritz, and R. R. Otter. 2008. The dark side of subsidies: adult stream insects export organic contaminants to riparian predators. Ecological Applications 18:1835–1841.

72. Wang, L., B. Ji, Y. Hu, R. Liu, and W. Sun. 2017. A review on in situ phytoremediation of mine tailings. Chemosphere 184:594–600.

73. Wilcove, D. S., D. Rothstein, J. Dubow, A. Phillips, and E. Losos. 1998. Quantifying threats to imperiled species in the United States. BioScience 48:607–615.

