## Supplemental Information 1 for "Integrating ecosystem and contaminant models to predict the effects of ecosystem fluxes on contaminant dynamics"

**Appendix S1**

Model Justification

While simple, the ecosystem model we use here is commonly used and captures basic biotic and abiotic feedbacks (e.g., consumption, excretions, mortality) in local ecosystems as well as some connections to neighbouring ecosystems. We use a wide range of parameter values (0 – 10) and a large number of parameter sets to explore general model behaviour. These values are always within the realm of reality, however, as we ensure we remove unfeasible solutions (e.g., an herbivore with a negative biomass cannot exist) and the mechanistic nature of the model ensures that other unfeasible solutions (e.g., an herbivore exists without autotroph prey items) never arise.

Individual parameter range justifications:

Input rate of inorganic (*I*): inorganic nutrient inputs for St Aubin’s Bay, New Jersey have been measured to range from 0 – 37 g day^−1^ for nitrogen, and lower inputs for phosphorus for the month of August depending on catchment and flow rate, thus our range of 0 – 10 is reasonable (Stapleton et al. 2000). We allow for *I* to be relatively large because previous work has shown how basal productivity impacts ecosystem dynamics (e.g., Oksanen et al. 1981).

Loss rate of inorganic nutrient (*l*): our range of 0 – 10 is appropriate by similar logic as *I* (input has to come from somewhere).

Assimilation efficiency of trophic level *i* (*e_i_*): is typically assumed to be 0.45 for consumption of primary producers and 0.85 for other feeding links (Yodzis and Innes 1992), however, see Cummins and Klug (1979) for a review of stream invertebrates estimating that assimilation efficiencies can be between 6 – 92% depending on food item and functional feeding guild of invertebrate and that some specific invertebrates can have even lower assimilation efficiencies (e.g., blackfly larvae; Wotton 1978). Thus, our assumed efficiency of 0 – 1 (proportion) is supported.

Consumption rate of trophic level *i* (*a_i_*): a review of stream invertebrate feeding measured consumption rates to be 30 – 50% of body weight / day, while fish consume 1 – 5 % of their body weight / day (Ng et al. 2000) which would mean that our consumption rate of 0 – 10 encompasses stream invertebrates up to 20 g, and fish up to 200 g in size.

Recycling rate of trophic level *i* (*d_i_*): our range of 0 – 10 time^−1^ is supported by a review by Vanni (2002) which reported nutrient excretion rates of freshwater invertebrates (10 – 85 mg_N_ m^−2^ d^−1^). Even considering the upper limit, 85 mg_N_ m^−2^ d^−1^, and taking the highest density of invertebrates (162 ind m^−2^) reported for a range of riffles in Wyoming (Hubert et al. 1996), approximating an average individual invertebrate biomass (e.g., chironomid) at 5 mg we would estimate *d_i_* to be 6.5 time^−1^ if the chironomid is completely made out of nutrient. Obviously it is not, and there will be a fraction of mortality over this time as well, so our range of 0 – 10 is supported.

Proportion of material lost from trophic level *i* (*δ_i_*): this is the proportion of material lost from a trophic level, e.g., via process of plant senescence or sedimentation of detritus. This is bounded by 0 and 1 because it is a proportion and values depend on species, temperature, time of year, etc. (Darchambeau et al. 2005).

Across ecosystem autotroph flux (*w_S_*): Our range of 0 – 10 is within the measured range (0 – 3 g m^−2^ day^−1^) of leaf litter to forested streams by Richardson et al. (2010).

**References:**

Cummins, K. W., and M. J. Klug. 1979. Feeding ecology of stream invertebrates. Annual review of ecology and systematics 10:147–172.

Darchambeau, F., I. Thys, B. Leporcq, L. Hoffmann, and J. Descy. 2005. Influence of zooplankton stoichiometry on nutrient sedimentation in a lake system. Limnology and Oceanography 50:905–913.

Hubert, W. A., W. J. LaVoie IV, and L. D. DeBray. 1996. Densities and substrate associations of macroinvertebrates in riffles of a small, high plains stream. Journal of Freshwater Ecology 11:21–26.

Ng, W.-K., K.-S. Lu, R. Hashim, and A. Ali. 2000. Effects of feeding rate on growth, feed utilizationand body composition of a tropical bagrid catfish. Aquaculture International 8:19–29.

Oksanen, L., S. D. Fretwell, J. Arruda, and P. Niemela. 1981. Exploitation ecosystems in gradients of primary productivity. The American Naturalist 118:240–261.

Richardson, J. S., Y. Zhang, and L. B. Marczak. 2010. Resource subsidies across the land–freshwater interface and responses in recipient communities. River Research and Applications 26:55–66.

Stapleton, C. M., D. Kay, G. F. Jackson, and M. D. Wyer. 2000. Estimated inorganic nutrient inputs to the coastal waters of Jersey from catchment and waste water sources. Water Research 34:787–796.

Vanni, M. J. 2002. Nutrient cycling by animals in freshwater ecosystems. Annual Review of Ecology and Systematics 33:341–370.

Wotton, R. 1978. Growth, respiration, and assimilation of blackfly larvae (Diptera: Simuliidae) in a lake-outlet in Finland. Oecologia 33:279–290.

Yodzis, P., and S. Innes. 1992. Body size and consumer-resource dynamics. The American Naturalist 139:1151–1175.
