## Supplemental Information 2 for "Integrating ecosystem and contaminant models to predict the effects of ecosystem fluxes on contaminant dynamics"

**Appendix S2: Base Model Equilibria**

Biomass Model:

Feasible Equilibrium:

$$N^{*}=\frac{a_{H}e_{H}I-d_{A}d_{H}(\delta_{A}-\delta_{H}e_{H}+e_{H}-1)}{a_{A}d_{H}((\delta_{H}-1)e_{H}+1)+a_{H}e_{H}l}$$

$$A^{*}=\frac{d_{H}}{a_{H}e_{H}}$$

$$H^{*}=\frac{a_{A}a_{H}e_{H}I-a_{A}\delta_{A}d_{A}d_{H}-a_{H}e_{H}ld_{A}}{a_{H}(a_{A}d_{H}((\delta_{H}-1)e_{H}+1)+a_{H}e_{H}l)}$$

Contaminant mass Model

$${C_{N}}^{*}=\frac{-a_{H}H^{*}(\delta_{H}-1)E_{1}(A^{*}fu_{A}+H^{*}u_{H})+a_{H}H^{*}E_{0}I+E_{1}d_{A}(A^{*}(u_{A}-\delta_{A}u_{A})-H^{*}(\delta_{H}-1)u_{H})+E_{0}Id_{A}}{a_{A}A^{*}(a_{H}H^{*}((\delta_{H}-1)f^{2}+1)+(\delta_{A}-1)fd_{A}+d_{A})+l(a_{H}H^{*}+d_{A})}$$

$${C_{A}}^{*}=\frac{A^{*}(a_{A}(A^{*}E_{1}u_{A}-H^{*}(\delta_{H}-1)E_{1}fu_{H}+E_{0}fI)+E_{1}lu_{A})}{a_{A}A^{*}(a_{H}H^{*}((\delta_{H}-1)f^{2}+1)+(\delta_{A}-1)fd_{A}+d_{A})+l(a_{H}H^{*}+d_{A})}$$

$${C_{H}}^{*}=\frac{H^{*}E_{1}u_{H}(a_{A}A^{*}(a_{H}H^{*}+(\delta_{A}-1)fd_{A}+d_{A})+l(a_{H}H^{*}+d_{A}))+a_{H}A^{*}H^{*}f(E_{1}u_{A}(a_{A}A^{*}+l)+a_{A}E_{0}fI)}{d_{H}(a_{A}A^{*}(a_{H}H^{*}((\delta_{H}-1)f^{2}+1)+(\delta_{A}-1)fd_{A}+d_{A})+l(a_{H}H^{*}+d_{A}))}$$

Concentrations:

$${\frac{C_{N}}{N}}^{*}=\frac{a_{H}H^{*}E_{0}I+E_{0}Id_{A}-a_{H}H^{*}(-1+\delta_{H})E_{1}(A^{*}fu_{A}+H^{*}u_{H})+E_{1}d_{A}(A^{*}(u_{A}-\delta_{A}u_{A})-H^{*}(-1+\delta_{H})u_{H})}{N^{*}l(a_{H}H^{*}+d_{A})+a_{A}N^{*}A^{*}(a_{H}H^{*}(1+(-1+\delta_{H})f^{2})+d_{A}+(-1+\delta_{A})fd_{A})}$$

$${\frac{C_{A}}{A}}^{*}=\frac{E_{1}lu_{A}+a_{A}(E_{0}fI+A^{*}E_{1}u_{A}-H^{*}(-1+\delta_{H})E_{1}fu_{H})}{l(a_{H}H^{*}+d_{A})+a_{A}A^{*}(a_{H}H^{*}(1+(-1+\delta_{H})f^{2})+d_{A}+(-1+\delta_{A})fd_{A})}$$

$${\frac{C_{H}}{H}}^{*}=\frac{a_{H}A^{*}f(a_{A}E_{0}fI+E_{1}(a_{A}A^{*}+l)u_{A})+E_{1}(l(a_{H}H^{*}+d_{A})+a_{A}A^{*}(a_{H}H^{*}+d_{A}+(-1+\delta_{A})fd_{A}))u_{H}}{(l(a_{H}H^{*}+d_{A})+a_{A}A^{*}(a_{H}H^{*}(1+(-1+\delta_{H})f^{2})+d_{A}+(-1+\delta_{A})fd_{A}))d_{H}}$$

Other Equilibria:

Case 1:

$$N^{*}=\frac{I}{l}$$

$$A^{*}=0$$

$$H^{*}=0$$

Contaminant Mass Model

$${C_{N}}^{*}=\frac{E_{0}I}{l}$$

$${C_{A}}^{*}=0$$

$${C_{H}}^{*}=0$$

Concentrations:

$${\frac{C_{N}}{N}}^{*}=E_{0}$$

$${\frac{C_{A}}{A}}^{*}=0$$

$${\frac{C_{H}}{H}}^{*}=0$$

Case 2:

$$N^{*}=\frac{d_{A}}{a_{A}}$$

$$A^{*}=\frac{a_{A}I-ld_{A}}{a_{A}\delta_{A}m_{A}}$$

$$H^{*}=0$$

Contaminant Mass Model

$${C_{N}}^{*}=\frac{a_{A}\delta_{A}E_{0}Id_{A}-(-1+\delta_{A})E_{1}(a_{A}I-ld_{A})u_{A}}{a_{A}(a_{A}(1+(-1+\delta_{A})f)I+(-1+\delta_{A}+f-\delta_{A}f)ld_{A})}$$

$${C_{A}}^{*}=0$$

$${C_{H}}^{*}=0$$

Concentrations:

$${\frac{C_{N}}{N}}^{*}=E_{0}$$

$${\frac{C_{A}}{A}}^{*}=0$$

$${\frac{C_{H}}{H}}^{*}=0$$
