## Supplemental Information 3 for "Integrating ecosystem and contaminant models to predict the effects of ecosystem fluxes on contaminant dynamics"

**Appendix S3: Substitutions for the partial derivatives presented in Fig. 2.**

**Biomass:**

$$p_{1}=a_{H}e_{H}$$

$$p_{2}=a_{A}e_{H}$$

$$q_{1}=a_{H}e_{H}l+a_{A}\left( 1+\left( -1+\delta_{h} \right)e_{H} \right)d_{H}$$

$$q_{2}=a_{H}e_{H}l+a_{A}\left( 1+\left( -1+\delta_{H} \right)e_{H} \right)d_{H}$$

$$r_{1}=-a_{H}e_{H}(a_{H}e_{H}I-\left( -1+\delta_{A}+e_{H}-\delta_{H}e_{H} \right)d_{A}d_{H})$$

$$r_{2}=-a_{A}e_{H}(a_{H}e_{H}I-\left( -1+\delta_{A}+e_{H}-\delta_{H}e_{H} \right)d_{A}d_{H})$$

$$s_{1}=-a_{A}^{2}\left( -1+e_{H} \right)\left( 1+\left( -1+2\delta_{H} \right)e_{H} \right)d_{H}^{2}$$

$$s_{2}=2a_{A}a_{H}e_{H}\left( 1+\left( -1+\delta_{H} \right)e_{H} \right)d_{H}$$

$$s_{3}=a_{H}^{2}e_{H}^{2}$$

$$s_{4}=a_{A}^{2}\left( 1+\left( -1+\delta_{H} \right)e_{H} \right)^{2}d_{H}^{2}$$

$$s_{5}=2a_{A}a_{H}e_{H}\left( 1+\left( -1+\delta_{H} \right)e_{H} \right)d_{H}$$

$$s_{6}=a_{H}^{2}e_{H}^{2}$$

**Contaminant Mass:**

$$c_{1}=a_{H}e_{H}d_{A}d_{H}(a_{H}^{3}E_{1}{e_{H}}^{3}\left( -1+\delta_{A}+f-\delta_{H}f \right)l^{3}u_{A}+a_{A}a_{H}^{2}e_{H}^{2}l^{2}(-E_{0}(-1+\delta_{A}+e_{H}-\delta_{H}e_{H})(1+(-1+\delta_{H})e_{H}+\delta_{A}(-1+f)+f(-1+f-\delta_{H}f))d_{A}d_{H}+E_{1}(-1+\delta_{A}+f-\delta_{H}f)(\left( 3+2\left( -1+\delta_{H} \right)e_{H} \right)d_{H}u_{A}+\left( -1+\delta_{H} \right)e_{H}fd_{A}u_{H}))+{a1}^{2}a2e_{H}ld_{H}(-E_{0}(-1+\delta_{A}+e_{H}-\delta_{H}e_{H})((1+(-1+\delta_{H})e_{H})(2+(-1+\delta_{H})e_{H}+f(-2+f-\delta_{H}f))+\delta_{A}(2(-1+f)-(-1+\delta_{H})(e_{H}-2e_{H}f+f^{2})))d_{A}d_{H}+E_{1}(-(1+(-1+\delta_{H})e_{H})(3+(-1+\delta_{H})e_{H})(1-\delta_{A}+(-1+\delta_{H})f)d_{H}u_{A}-(-1+\delta_{H})e_{H}(\left( -1+\delta_{A}+e_{H}-\delta_{H}e_{H} \right)^{2}-2\left( -1+\delta_{A} \right)\delta_{A}f+2\delta_{A}(-1+\delta_{H})f^{2})d_{A}u_{H}))-a_{A}^{3}{d_{H}}^{2}(-E_{0}(1+(-1+\delta_{H})e_{H})(1-\delta_{A}+(-1+\delta_{H})e_{H})(-(1+(-1+\delta_{H})e_{H})(-1+f)+\delta_{A}(-1+f(1+(-1+\delta_{H})e_{H}+f-\delta_{H}f)))d_{A}d_{H}+E_{1}(-\left( 1+\left( -1+\delta_{H} \right)e_{H} \right)^{2}(-1+\delta_{A}+f-\delta_{H}f)d_{H}u_{A}+\left( -1+\delta_{H} \right)e_{H}\left( \left( -1+\delta_{A}+e_{H}-\delta_{H}e_{H} \right)^{2}-\left( -1+\delta_{A} \right)\left( 1+\left( -1+\delta_{H} \right)e_{H} \right)\left( -1+2\delta_{A}+e_{H}-\delta_{H}e_{H} \right)f+{\delta_{A}}^{2}\left( -1+\delta_{H} \right)f^{2} \right)d_{A}u_{H})))$$

$$c_{2}= 2a_{A}a_{H}^{2}{e_{H}}^{2}d_{A}d_{H}(a_{H}e_{H}(1+\left( -1+\delta_{H} \right)e_{H}+\delta_{A}(-1+f)+f(-1+f-\delta_{H}f))l+a_{A}(-(1+(-1+\delta_{H})e_{H})(-1+f)+\delta_{A}(-1+f(1+(-1+\delta_{H})e_{H}+f-\delta_{H}f)))d_{H})(a_{H}E_{0}e_{H}l+a_{A}E_{0}(1+(-1+\delta_{H})e_{H})d_{H}-a_{A}\left( -1+\delta_{H} \right)E_{1}e_{H}u_{H})$$

$$c_{3}= a_{A}{a_{H}}^{3}e_{H}^{3}(a_{H}e_{H}l+a_{A}(1+(-1+\delta_{H})f^{2})d_{H})(a_{H}E_{0}e_{H}l+a1E_{0}(1+(-1+\delta_{H})e_{H})d_{H}-a_{A}\left( -1+\delta_{H} \right)E_{1}e_{H}u_{H})$$

$$c_{4}= d_{H}(a_{H}e_{H}l+a_{A}(1+(-1+\delta_{H})e_{H})d_{H})({a_{H}}^{2}E_{1}{e_{H}}^{2}l^{2}u_{A}+a_{A}a_{H}e_{H}l(E_{0}f(-1+\delta_{A}+e_{H}-\delta_{H}e_{H}+f-\delta_{A}f+(-1+\delta_{H})f^{2})d_{A}d_{H}+E_{1}\left( 2+\left( -1+\delta_{H} \right)f^{2} \right)d_{H}u_{A}+(-1+\delta_{H})E_{1}e_{H}fd_{A}u_{H})+{a_{A}}^{2}d_{H}(E_{0}f(-1+\delta_{A}+e_{H}-\delta_{H}e_{H}-(-1+\delta_{A})(1+(-1+\delta_{H})e_{H})f+\delta_{A}(-1+\delta_{H})f^{2})d_{A}d_{H}+E_{1}(1+(-1+\delta_{H})f^{2})d_{H}u_{A}+(-1+\delta_{H})E_{1}e_{H}f(1+(-1+\delta_{A})f)d_{A}u2))$$

$$c_{5}= a_{A}d_{A}d_{H}(a_{H}^{4}E_{1}{e_{H}}^{4}fl^{4}u_{A}+a_{A}a_{H}^{3}{e_{H}}^{3}fl^{3}(E_{0}f\left( -1+\delta_{A}+e_{H}-\delta_{H}e_{H}+f-\delta_{A}f+\left( -1+\delta_{H} \right)f^{2} \right)d_{A}d_{H}+E_{1}\left( 4+2\left( -1+\delta_{H} \right)e_{H}+\left( -1+\delta_{A} \right)f \right)d_{H}u_{A}+\left( -1+\delta_{H} \right)E_{1}e_{H}fd_{A}u_{H})+{a_{A}}^{2}a_{H}^{2}{e2}^{2}l^{2}d_{H}(E_{0}f^{2}\left( \delta_{A}-{\delta_{A}}^{2}\left( -1+f \right)+\delta_{A}\left( -1-2\left( -1+\delta_{H} \right)e_{H} \right)f+2\delta_{A}\left( -1+\delta_{H} \right)f^{2}-\left( 1+\left( -1+\delta_{H} \right)e_{H} \right)\left( 2+\left( -1+\delta_{H} \right)e_{H}+f\left( -2+f-\delta_{H}f \right) \right) \right)d_{A}d_{H}+E_{1}f\left( 6+3\left( -1+\delta_{A} \right)f+\left( -1+\delta_{H} \right)e_{H}\left( 6+\left( -1+\delta_{H} \right)e_{H}+2\left( -1+\delta_{A} \right)f \right) \right)d_{H}u_{A}+E_{1}e_{H}\left( \left( -1+\delta_{A}+e_{H}-\delta_{H}e_{H} \right)^{2}-2\left( -1+\delta_{A} \right)\left( -1+\delta_{A}+e_{H}-\delta_{H}e_{H} \right)f+\left( \delta_{H}+\delta_{A}\left( -4+\delta_{A}+2\delta_{H} \right) \right)f^{2} \right)d_{A}u_{H})+{a1}^{4}{d_{H}}^{3}({\delta_{A}}^{2}d_{A}(-E_{0}\left( 1+\left( -1+\delta_{H} \right)e_{H} \right)f^{2}\left( -1+f\left( 1+\left( -1+\delta_{H} \right)e_{H}+f-\delta_{H}f \right) \right)d_{H}+E_{1}e_{H}\left( 1+f\left( -2+2\left( -1+\delta_{H} \right)e_{H}\left( -1+f \right)+\delta_{H}f+\left( -1+\delta_{H} \right)^{2}{e_{H}}^{2}f \right) \right)u_{H})+E_{1}\left( 1+\left( -1+\delta_{H} \right)e_{H} \right)^{2}(-1+f)(-fd_{H}u_{A}+e_{H}\left( -1+f \right)d_{A}u_{H})+\delta_{A}(1+(-1+\delta_{H})e_{H})(\left( 1+\left( -1+\delta_{H} \right)e_{H} \right)f^{2}d_{H}(E_{0}\left( -1+f \right)d_{A}+E_{1}u_{A})-2E_{1}e_{H}\left( -1+f \right)\left( -1+f+\left( -1+\delta_{H} \right)e_{H}f \right)d_{A}u_{H}))+{a_{A}}^{3}a_{H}e_{H}l{d_{H}}^{2}(E_{0}f^{2}\left( \left( 1+\left( -1+\delta_{H} \right)e_{H} \right)^{2}\left( -1+f \right)+\delta_{A}\left( 1+\left( -1+\delta_{H} \right)e_{H} \right)\left( -1+f-\left( -1+\delta_{H} \right)\left( e_{H}+e_{H}f-2f^{2} \right) \right)+{\delta_{A}}^{2}\left( 2-2f+\left( -1+\delta_{H} \right)\left( e_{H}-2e_{H}f+f^{2} \right) \right) \right)d_{A}d_{H}+E_{1}(2e_{H}\left( -1+\delta_{A}+e_{H}-\delta_{H}e_{H} \right)^{2}d_{A}u_{H}+f^{2}(\left( -1+\delta_{A} \right)\left( 1+\left( -1+\delta_{H} \right)e_{H} \right)\left( 3+\left( -1+\delta_{H} \right)e_{H} \right)d_{H}u_{A}+e_{H}\left( 2-6\delta_{A}+{\delta_{A}}^{2}+2\delta_{A}\delta_{H}+{\delta_{A}}^{2}\delta_{H}+2\left( -1+\delta_{A} \right)^{2}\left( -1+\delta_{H} \right)e_{H} \right)d_{A}u_{H})+2(2+(-1+\delta_{H})e_{H})f(\left( 1+\left( -1+\delta_{H} \right)e_{H} \right)d_{H}u_{A}-\left( -1+\delta_{A} \right)e_{H}\left( -1+\delta_{A}+e_{H}-\delta_{H}e_{H} \right)d_{A}u_{H}))))$$

$$c_{6}= 2{a_{A}}^{3}a_{H}e_{H}d_{A}d_{H}\left( a_{H}e_{H}\left( 1+\left( -1+\delta_{H} \right)e_{H}+\delta_{A}\left( -1+f \right)+f\left( -1+f-\delta_{H}f \right) \right)l+a_{A}\left( -\left( 1+\left( -1+\delta_{H} \right)e_{H} \right)\left( -1+f \right)+\delta_{A}\left( -1+f\left( 1+\left( -1+\delta_{H} \right)e_{H}+f-\delta_{H}f \right) \right) \right)d_{H} \right)\left( a_{H}e_{H}l\left( E_{0}f^{2}d_{H}+E_{1}e_{H}u_{H} \right)+a_{A}d_{H}\left( E_{0}\left( 1+\left( -1+\delta_{H} \right)e_{H} \right)f^{2}d_{H}+E_{1}e_{H}u_{H} \right) \right)$$

$${c_{7}= a_{A}}^{3}a_{H}^{2}{e_{H}}^{2}(a_{H}e_{H}l+a_{A}\left( 1+\left( -1+\delta_{H} \right)f^{2} \right)d_{H})(a_{H}e_{H}l(E_{0}f^{2}d_{H}+E_{1}e_{H}u_{H})+a_{A}d_{H}(E_{0}\left( 1+\left( -1+\delta_{H} \right)e_{H} \right)f^{2}d_{H}+E_{1}e_{H}u_{H}))$$

$$g_{1}= a_{A}{d_{A}}^{2}{d_{H}}^{2}(a_{H}e_{H}l+a_{A}(1+(-1+\delta_{H})e_{H})d_{H}){(a_{H}e_{H}(1+\left( -1+\delta_{H} \right)e_{H}+\delta_{A}(-1+f)+f(-1+f-\delta_{H}f))l+a_{A}(-(1+(-1+\delta_{H})e_{H})(-1+f)+\delta_{A}(-1+f(1+(-1+\delta_{H})e_{H}+f-\delta_{H}f)))d_{H})}^{2}$$

$$g_{2}= 2a_{A}a_{H}e_{H}d_{A}d_{H}(a_{H}e_{H}l+a_{A}(1+(-1+\delta_{H})e_{H})d_{H})(a_{H}e_{H}l+a_{A}(1+(-1+\delta_{H})f^{2})d_{H})(a_{H}e_{H}(1+(-1+\delta_{H})e2+\delta_{A}(-1+f)+f(-1+f-\delta_{H}f))l+a_{A}(-(1+(-1+\delta_{H})e_{H})(-1+f)+\delta_{A}(-1+f(1+(-1+\delta_{H})e_{H}+f-\delta_{H}f)))d_{H})$$

$$g_{3}= a_{A}{a_{H}}^{2}{e_{H}}^{2}(a_{H}e_{H}l+a_{A}(1+(-1+\delta_{H})e_{H})d_{H}){(a_{H}e_{H}l+a_{A}(1+(-1+\delta_{H})f^{2})d_{H})}^{2}$$

$$g_{4}=a_{A}{d_{A}}^{2}{d_{H}}^{2}{(a_{H}e_{H}(1+\left( -1+d_{H} \right)e_{H}+\delta_{A}(-1+f)+f(-1+f-\delta_{H}f))l+a_{A}(-(1+(-1+\delta_{H})e_{H})(-1+f)+\delta_{A}(-1+f(1+(-1+\delta_{H})e_{H}+f-\delta_{H}f)))d_{H})}^{2}$$

$$g_{5}= 2a_{A}a_{H}e_{H}d_{A}d_{H}(a_{H}e_{H}l+a_{A}(1+(-1+\delta_{H})f^{2})d_{H})(a_{H}e_{H}(1+(-1+\delta_{H})e_{H}+\delta_{A}(-1+f)+f(-1+f-\delta_{H}f))l+a_{A}(-(1+(-1+\delta_{H})e_{H})(-1+f)+\delta_{A}(-1+f(1+(-1+\delta_{H})e_{H}+f-\delta_{H}f)))d_{H})$$

$$g_{6}= a_{A}a_{H}^{2}e_{H}^{2}{(a_{H}e_{H}l+a_{A}\left( 1+\left( -1+\delta_{H} \right)f^{2} \right)d_{H})}^{2}$$

$$g_{7}={a_{A}}^{2}d_{H}(a_{H}e_{H}l+a_{A}\left( 1+\left( -1+\delta_{H} \right)e_{H} \right)d_{H})$$

$$g_{8}= a_{H}^{2}{e_{H}}^{2}l+a_{H}e_{H}(a_{A}(1+(-1+\delta_{H})f^{2})$$

$$g_{9}= \left( 1+\left( -1+\delta_{H} \right)e_{H}+\delta_{A}\left( -1+f \right)+f\left( -1+f-\delta_{H}f \right) \right)ld_{A})d_{H} +a_{A}(-(1+(-1+\delta_{H})e_{H})(-1+f)+\delta_{A}(-1+f(1+(-1+\delta_{H})e_{H}+f-\delta_{H}f)))d_{A} d_{H}^{2}$$

$$k_{1}=-a_{A}^{4}d_{H}\left( a_{H}e_{H}I-\left( -1+\delta_{A}+e_{H}-\delta_{H}e_{H} \right)d_{A}d_{H} \right)\left( a_{H}^{2}e_{H}^{2}I^{2}\left( E_{0}\left( 1+\left( -1+\delta_{H} \right)e_{H} \right)^{2}d_{H}-\left( -1+\delta_{H} \right)E_{1}e_{H}\left( 2+\left( -1+\delta_{H} \right)e_{H}+\left( -1+\delta_{H} \right)f^{2} \right)u_{H} \right)-\left( -1+\delta_{H} \right)E_{1}d_{A}{d_{H}}^{2}\left( -\left( -1+\delta_{A} \right)fd_{H}u_{A}-\left( -1+\delta_{A} \right)e_{H}\left( -1+2\left( -1+\delta_{H} \right)f \right)d_{A}u_{H}+e_{H}\left( 1+3\delta_{A}\left( -1+f \right)-f+{\delta_{A}}^{2}\left( 2+f\left( -2+\left( -1+\delta_{H} \right)f \right) \right) \right)d_{A}u_{H}+\left( -1+\delta_{A} \right)\left( -1+\delta_{H} \right)^{2}e_{H}^{3}\left( d_{H}u_{A}+\left( -1+f \right)d_{A}u_{H} \right)+\left( -1+\delta_{H} \right){e_{H}}^{2}\left( -\left( -1+\delta_{A} \right)\left( -2+\left( -1+\delta_{H} \right)f \right)d_{H}u_{A}+\left( 2-4\delta_{A}+{\delta_{A}}^{2}-2\left( -1+\delta_{A} \right)^{2}f \right)d_{A}u_{H} \right) \right)+a_{H}e_{H}Id_{H}\left( E_{0}\left( 1+\left( -1+\delta_{H} \right)e_{H} \right)^{2}\left( 1+\left( -1+\delta_{H} \right)e_{H}+\delta_{A}\left( -1+f \right)+f\left( -1+f-\delta_{H}f \right) \right)d_{A}d_{H}-\left( -1+\delta_{H} \right)E_{1}\left( \left( 1+\left( -1+\delta_{H} \right)e_{H} \right)^{2}fd_{H}u_{A}+e_{H}\left( \left( 1+\left( -1+\delta_{H} \right)e_{H} \right)\left( 3+\left( -1+\delta_{H} \right)e_{H}-2f \right)+2\delta_{A}\left( -2+f+\left( -1+\delta_{H} \right)\left( e_{H}\left( -1+f \right)-f^{2} \right) \right) \right)d_{A}u_{H} \right) \right) \right)$$

$$k_{2}= -2a_{A}^{3}a_{H}e_{H}(a_{H}e_{H}I-\left( -1+\delta_{A}+e_{H}-\delta_{H}e_{H} \right)d_{A}d_{H})(a_{H}^{2}e_{H}^{2}I^{2}(E_{0}\left( 1+\left( -1+\delta_{H} \right)e_{H} \right)d_{H}-\left( -1+\delta_{H} \right)E_{1}e_{H}u_{H})-\left( -1+\delta_{H} \right)E_{1}d_{A}{d_{H}}^{2}(\left( -1+\delta_{A} \right)\left( 1+\left( -1+\delta_{H} \right)e_{H} \right)\left( e_{H}-f \right)d_{H}u_{A}+\delta_{A}e_{H}\left( -1+\delta_{A}+e_{H}-\delta_{H}e_{H}+f-\delta_{A}f+\left( -1+\delta_{H} \right)f^{2} \right)d_{A}u_{H})+a_{H}e_{H}Id_{H}(E_{0}\left( 1+\left( -1+\delta_{H} \right)e_{H} \right)\left( 1+\left( -1+\delta_{H} \right)e_{H}+\delta_{A}\left( -1+f \right)+f\left( -1+f-\delta_{H}f \right) \right)d_{A}d_{H}+\left( -1+\delta_{H} \right)E_{1}(\left( -1+e_{H}-\delta_{H}e_{H} \right)fd_{H}u_{A}+e_{H}\left( -1+e_{H}-\delta_{H}e_{H}-\delta_{A}\left( -2+f \right)+f+\left( -1+\delta_{H} \right)f^{2} \right)d_{A}u_{H})))$$

$$k_{3}= -a_{A}^{2}a_{H}^{2}e_{H}^{2}(a_{H}e_{H}I-\left( -1+\delta_{A}+e_{H}-\delta_{H}e_{H} \right)d_{A}d_{H})(a_{H}^{2}E_{0}e_{H}^{2}I^{2}+\left( -1+\delta_{H} \right)E_{1}d_{A}d_{H}(\left( -1+\delta_{A} \right)\left( -e_{H}+f \right)d_{H}u_{A}+e_{H}\left( 1+\left( -1+\delta_{H} \right)e_{H}+\delta_{A}\left( -1+f \right)+f\left( -1+f-\delta_{H}f \right) \right)d_{A}u_{H})+a_{H}e_{H}I(E_{0}\left( 1+\left( -1+\delta_{H} \right)e_{H}+\delta_{A}\left( -1+f \right)+f\left( -1+f-\delta_{H}f \right) \right)d_{A}d_{H}-\left( -1+\delta_{H} \right)E_{1}(fd_{H}u_{A}-e_{H}d_{A}u_{H})))$$

$$m_{1}=a_{A}^{2}d_{H}^{2}{(a_{H}e_{H}(1+(-1+\delta_{H})f^{2})I+\left( -\left( 1+\left( -1+\delta_{H} \right)e_{H} \right)\left( -1+f \right)+\delta_{A}\left( -1+f\left( 1+\left( -1+\delta_{H} \right)e_{H}+f-\delta_{H}f \right) \right) \right)d_{A}d_{H})}^{2}$$

$$m_{2}=2a_{A}a_{H}e_{H}d_{H}(a_{H}e_{H}I+\left( 1+\left( -1+\delta_{H} \right)e_{H}+\delta_{A}\left( -1+f \right)+f\left( -1+f-\delta_{H}f \right) \right)d_{A}d_{H})(a_{H}e_{H}(1+(-1+\delta_{H})f^{2})I+\left( -\left( 1+\left( -1+\delta_{H} \right)e_{H} \right)\left( -1+f \right)+\delta_{A}\left( -1+f\left( 1+\left( -1+\delta_{H} \right)e_{H}+f-\delta_{H}f \right) \right) \right)d_{A}d_{H})$$

$$m_{3}=a_{H}^{2}e_{H}^{2}{(a_{H}e_{H}I+\left( 1+\left( -1+\delta_{H} \right)e_{H}+\delta_{A}\left( -1+f \right)+f\left( -1+f-\delta_{H}f \right) \right)d_{A}d_{H})}^{2}$$

$$m_{4}=a_{A}^{4}{(d_{H}+\left( -1+\delta_{H} \right)e_{H}d_{H})}^{2}$$

$$m_{5}= 2a_{A}^{3}a_{H}e_{H}\left( 1+\left( -1+\delta_{H} \right)e_{H} \right)d_{H}$$

$$m_{6}= a_{A}^{2}a_{H}^{2}e_{H}^{2}$$

$$k_{4}= -{a_{A}}^{4}(1+\left( -1+\delta_{H} \right)e_{H})d_{H}^{2}(a_{H}^{2}(-1+\delta_{H})e_{H}^{2}fI^{2}(E_{0}\left( e_{H}-f^{2} \right)d_{H}-E_{1}e_{H}u_{H})+E_{1}d_{A}{d_{H}}^{2}(\left( -1+\delta_{A}+e_{H}-\delta_{H}e_{H}-\left( -1+\delta_{A} \right)\left( 1+\left( -1+\delta_{H} \right)e_{H} \right)^{2}f+\left( -1+\delta_{H} \right)\left( -1+2\delta_{A}+\left( -1+\delta_{A} \right)\left( -1+\delta_{H} \right)e_{H} \right)f^{2} \right)d_{H}u_{A}+\left( -1+\delta_{A} \right)\left( -1+\delta_{H} \right)e_{H}\left( -1+\delta_{A}+e_{H}-\delta_{H}e_{H} \right)\left( -1+f \right)fd_{A}u_{H})+a_{H}e_{H}Id_{H}(\left( -1+\delta_{H} \right)E_{0}\left( 1-\delta_{A}+\left( -1+\delta_{H} \right)e_{H} \right)f\left( e_{H}-f^{2} \right)d_{A}d_{H}-E_{1}(\left( 1+\left( -1+\delta_{H} \right)\left( 2+\left( -1+\delta_{H} \right)e_{H} \right)f^{2} \right)d_{H}u_{A}+\left( -1+\delta_{H} \right)e_{H}\left( 2+\left( -1+\delta_{H} \right)e_{H}+\delta_{A}\left( -2+f \right)-f \right)fd_{A}u_{H})))$$

$$k_{5}= a_{A}^{3}a_{H}e_{H}d_{H}(-{a_{H}}^{2}(-1+\delta_{H}){e_{H}}^{2}fI^{2}(E_{0}\left( e_{H}-f^{2} \right)d_{H}-E_{1}e_{H}u_{H})+a_{H}e_{H}Id_{H}(-\left( -1+\delta_{H} \right)E_{0}\left( 1-\delta_{A}+\left( -1+\delta_{H} \right)e_{H} \right)f\left( e_{H}-f^{2} \right)d_{A}d_{H}+E_{1}\left( 3-2e_{H}+2\delta_{H}e_{H}+\left( -1+\delta_{H} \right)\left( 4+3\left( -1+\delta_{H} \right)e_{H} \right)f^{2} \right)d_{H}u_{A}+\left( -1+\delta_{H} \right)E_{1}e_{H}\left( 2+\left( -1+\delta_{H} \right)e_{H}+\delta_{A}\left( -2+f \right)-f \right)fd_{A}u_{H})+E_{1}d_{A}{d_{H}}^{2}(\left( -\left( 3+2\left( -1+\delta_{H} \right)e_{H} \right)\left( -1+\delta_{A}+e_{H}-\delta_{H}e_{H} \right)+3\left( -1+\delta_{A} \right)\left( 1+\left( -1+\delta_{H} \right)e_{H} \right)^{2}f-\left( -1+\delta_{H} \right)\left( -1+4\delta_{A}+\left( -1+3\delta_{A} \right)\left( -1+\delta_{H} \right)e_{H} \right)f^{2} \right)d_{H}u_{A}-\left( -1+\delta_{A} \right)\left( -1+\delta_{H} \right)e_{H}\left( -1+\delta_{A}+e_{H}-\delta_{H}e_{H} \right)\left( -1+f \right)fd_{A}u_{H}))$$

$$k_{6}= a_{A}^{2}a_{H}^{2}E_{1}e_{H}^{2}d_{H}^{2}\left( a_{H}e_{H}\left( 3+\left( -1+\delta_{H} \right)e_{H}+2\left( -1+\delta_{H} \right)f^{2} \right)J+\left( \left( 1+\left( -1+\delta_{H} \right)e_{H} \right)\left( 3+\left( -1+\delta_{H} \right)e_{H}+f\left( -3+f-\delta_{H}f \right) \right)+\delta_{A}\left( 3\left( -1+f \right)-\left( -1+\delta_{H} \right)\left( e_{H}-3e_{H}f+2f^{2} \right) \right) \right)d_{A}d_{H} \right)u_{A}$$

$$k_{7}= a_{A}{a_{H}}^{3}E_{1}{e_{H}}^{3}d_{H}\left( a_{H}e_{H}I+\left( 1+\left( -1+\delta_{H} \right)e_{H}+\delta_{A}\left( -1+f \right)+f\left( -1+f-\delta_{H}f \right) \right)d_{A}d_{H} \right)u_{A}$$

$$k_{8}= -{a_{A}}^{5}{d_{H}}^{2}({a_{H}}^{3}{e_{H}}^{3}I^{3}(E_{0}\left( f+\left( -1+\delta_{H} \right)e_{Hf} \right)^{2}d_{H}-E_{1}e_{H}\left( -1+\left( -1+\delta_{H} \right)^{2}e_{H}f^{2} \right)u_{H})+{a_{H}}^{2}{e_{H}}^{2}I^{2}d_{H}(E_{0}\left( 2+\left( -1+\delta_{H} \right)e_{H}+\delta_{A}\left( -2+f \right)-f \right)\left( f+\left( -1+\delta_{H} \right)e_{Hf} \right)^{2}d_{A}d_{H}-\left( -1+\delta_{H} \right)E_{1}\left( 1+\left( -1+\delta_{H} \right)e_{H} \right)^{2}f^{3}d_{H}u_{A}-E_{1}e_{H}\left( 3\left( -1+\delta_{A}+e_{H}-\delta_{H}e_{H} \right)-2\left( -1+\delta_{A} \right)\left( 1+\left( -1+\delta_{H} \right)e_{H} \right)f+\left( -1+\delta_{H} \right)^{2}e_{H}\left( 1-3\delta_{A}+\left( -1+\delta_{H} \right)e_{H} \right)f^{2} \right)d_{A}u_{H})-E_{1}{d_{A}}^{2}{d_{H}}^{3}(\left( -1+\delta_{H} \right)\left( \delta_{A}+\delta_{A}\left( -1+\delta_{H} \right)e_{H} \right)^{2}f^{3}d_{H}u_{A}+e_{H}\left( -1+\delta_{A}+e_{H}-\delta_{H}e_{H} \right)^{3}d_{A}u_{H}+(1+\left( -1+\delta_{H} \right)e_{H})(-1+\delta_{A}+e_{H}-\delta_{H}e_{H})f(\left( 1+\left( -1+\delta_{H} \right)e_{H} \right)d_{H}u_{A}-2\left( -1+\delta_{A} \right)e_{H}\left( -1+\delta_{A}+e_{H}-\delta_{H}e_{H} \right)d_{A}u_{H})+f^{2}(-\left( -1+\delta_{A} \right)\left( 1+\left( -1+\delta_{H} \right)e_{H} \right)^{2}\left( 1+\left( 1+\delta_{A} \right)\left( -1+\delta_{H} \right)e_{H} \right)d_{H}u_{A}-e_{H}\left( 1-\delta_{A}+\left( -1+\delta_{H} \right)e_{H} \right)\left( \left( -1+\delta_{A} \right)^{2}-\left( -2+\delta_{A}\left( 4+\delta_{A}\left( -3+\delta_{H} \right) \right) \right)\left( -1+\delta_{H} \right)e_{H}+\left( -1+\delta_{A} \right)^{2}\left( -1+\delta_{H} \right)^{2}{e_{H}}^{2} \right)d_{A}u_{H}))+a_{H}e_{H}Id_{A}{d_{H}}^{2}(-\left( -1+\delta_{A} \right)E_{0}\left( -1+\delta_{A}+e_{H}-\delta_{H}e_{H} \right)\left( -1+f \right)\left( f+\left( -1+\delta_{H} \right)e_{Hf} \right)^{2}d_{A}d_{H}+E_{1}(2\delta_{A}\left( -1+\delta_{H} \right)\left( 1+\left( -1+\delta_{H} \right)e_{H} \right)^{2}f^{3}d_{H}u_{A}+3e_{H}\left( -1+\delta_{A}+e_{H}-\delta_{H}e_{H} \right)^{2}d_{A}u_{H}+(1+\left( -1+\delta_{H} \right)e_{H})f(\left( 1+\left( -1+\delta_{H} \right)e_{H} \right)d_{H}u_{A}-4\left( -1+\delta_{A} \right)e_{H}\left( -1+\delta_{A}+e_{H}-\delta_{H}e_{H} \right)d_{A}u_{H})+e_{H}f^{2}(-\left( -1+\delta_{A} \right)\left( -1+\delta_{H} \right)\left( 1+\left( -1+\delta_{H} \right)e_{H} \right)^{2}d_{H}u_{A}+\left( \left( -1+\delta_{A} \right)^{2}-\left( -1+\delta_{H} \right)\left( -2+\delta_{A}\left( 6-5\delta_{A}-2\delta_{H}+3\delta_{A}\delta_{H} \right) \right)e_{H}+\left( -1+\delta_{H} \right)^{2}\left( 1+\delta_{A}\left( -4+\delta_{A}+2\delta_{H} \right) \right){e_{H}}^{2} \right)d_{A}u_{H}))))$$

$$k_{9}= -2{a_{A}}^{4}a_{H}e_{H}d_{H}({a_{H}}^{3}{e_{H}}^{3}I^{3}(E_{0}\left( 1+\left( -1+\delta_{H} \right)e_{H} \right)f^{2}d_{H}+E_{1}e_{H}u_{H})+{a_{H}}^{2}{e_{H}}^{2}I^{2}d_{H}(E_{0}\left( 1+\left( -1+\delta_{H} \right)e_{H} \right)\left( 2+\left( -1+\delta_{H} \right)e_{H}+\delta_{A}\left( -2+f \right)-f \right)f^{2}d_{A}d_{H}-\left( -1+\delta_{H} \right)E_{1}\left( 1+\left( -1+\delta_{H} \right)e_{H} \right)f^{3}d_{H}u_{A}+E_{1}e_{H}\left( -3\left( -1+\delta_{A}+e_{H}-\delta_{H}e_{H} \right)+\left( -1+\delta_{A} \right)\left( 2+\left( -1+\delta_{H} \right)e_{H} \right)f+\left( -1+\delta_{H} \right)^{2}e_{H}f^{2} \right)d_{A}u_{H})+E_{1}{d_{A}}^{2}{d_{H}}^{3}(-\delta_{A}\left( -1+\delta_{H} \right)\left( 1+\left( -1+\delta_{H} \right)e_{H} \right)\left( 1+\delta_{A}+\left( -1+\delta_{H} \right)e_{H} \right)f^{3}d_{H}u_{A}+e_{H}\left( 1-\delta_{A}+\left( -1+\delta_{H} \right)e_{H} \right)^{3}d_{A}u_{H}+f^{2}(\left( -1+\delta_{A} \right)\left( 1+\left( -1+\delta_{H} \right)e_{H} \right)\left( 2+\left( -1+\delta_{H} \right)e_{H}\left( 3+\delta_{A}+\left( -1+\delta_{H} \right)e_{H} \right) \right)d_{H}u_{A}+e_{H}\left( 1-\delta_{A}+\left( -1+\delta_{H} \right)e_{H} \right)\left( \left( -1+\delta_{A} \right)^{2}+\left( 1+\delta_{A}\left( -1+\delta_{A}-\delta_{H} \right) \right)\left( -1+\delta_{H} \right)e_{H} \right)d_{A}u_{H})+(2+\left( -1+\delta_{H} \right)e_{H})(-1+\delta_{A}+e_{H}-\delta_{H}e_{H})f(\left( -1+e_{H}-\delta_{H}e_{H} \right)d_{H}u_{A}+\left( -1+\delta_{A} \right)e_{H}\left( -1+\delta_{A}+e_{H}-\delta_{H}e_{H} \right)d_{A}u_{H}))+a_{H}e_{H}Id_{A}{d_{H}}^{2}(-\left( -1+\delta_{A} \right)E_{0}\left( 1+\left( -1+\delta_{H} \right)e_{H} \right)\left( -1+\delta_{A}+e_{H}-\delta_{H}e_{H} \right)\left( -1+f \right)f^{2}d_{A}d_{H}+E_{1}(\left( -1+\delta_{H} \right)\left( 1+\left( -1+\delta_{H} \right)e_{H} \right)\left( 1+2\delta_{A}+\left( -1+\delta_{H} \right)e_{H} \right)f^{3}d_{H}u_{A}+3e_{H}\left( -1+\delta_{A}+e_{H}-\delta_{H}e_{H} \right)^{2}d_{A}u_{H}+(2+\left( -1+\delta_{H} \right)e_{H})f(\left( 1+\left( -1+\delta_{H} \right)e_{H} \right)d_{H}u_{A}-2\left( -1+\delta_{A} \right)e_{H}\left( -1+\delta_{A}+e_{H}-\delta_{H}e_{H} \right)d_{A}u_{H})+e_{H}f^{2}(-\left( -1+\delta_{A} \right)\left( -1+\delta_{H} \right)\left( 1+\left( -1+\delta_{H} \right)e_{H} \right)d_{H}u_{A}+\left( \left( -1+\delta_{A} \right)^{2}+\left( -1+\delta_{H} \right)\left( {\delta_{A}}^{2}+\delta_{H}-2\delta_{A}\delta_{H} \right)e_{H}+\left( -1+\delta_{H} \right)^{3}{e_{H}}^{2} \right)d_{A}u_{H}))))$$

$$k_{10}=-{a_{A}}^{3}{a_{H}}^{2}{e_{H}}^{2}({a_{H}}^{3}{e_{H}}^{3}I^{3}(E_{0}f^{2}d_{H}+E_{1}e_{H}u_{H})+a_{H}e_{H}Id_{A}{d_{H}}^{2}(-\left( -1+\delta_{A} \right)E_{0}\left( -1+\delta_{A}+e_{H}-\delta_{H}e_{H} \right)\left( -1+f \right)f^{2}d_{A}d_{H}+E_{1}f\left( 6+\left( -1+\delta_{H} \right)\left( e_{H}\left( 6+\left( -1+\delta_{H} \right)e_{H} \right)-\left( -1+\delta_{A} \right)e_{H}f+2\left( 2+\delta_{A}+2\left( -1+\delta_{H} \right)e_{H} \right)f^{2} \right) \right)d_{H}u_{A}+E_{1}e_{H}\left( 3\left( -1+\delta_{A}+e_{H}-\delta_{H}e_{H} \right)^{2}-4\left( -1+\delta_{A} \right)\left( -1+\delta_{A}+e_{H}-\delta_{H}e_{H} \right)f+\left( \left( -1+\delta_{A} \right)^{2}-\left( -1+\delta_{H} \right)^{2}e_{H} \right)f^{2} \right)d_{A}u_{H})+{a_{H}}^{2}{e_{H}}^{2}I^{2}d_{H}(f^{2}d_{H}(E_{0}\left( 2+\left( -1+\delta_{H} \right)e_{H}+\delta_{A}\left( -2+f \right)-f \right)d_{A}-\left( -1+\delta_{H} \right)E_{1}fu_{A})+E_{1}e_{H}\left( 3+3\left( -1+\delta_{H} \right)e_{H}-2f+\delta_{A}\left( -3+2f \right) \right)d_{A}u_{H})-E_{1}{d_{A}}^{2}{d_{H}}^{3}(\left( -1+\delta_{H} \right)\left( {\delta_{A}}^{2}+4\delta_{A}\left( 1+\left( -1+\delta_{H} \right)e_{H} \right)+\left( 1+\left( -1+\delta_{H} \right)e_{H} \right)^{2} \right)f^{3}d_{H}u_{A}+e_{H}\left( -1+\delta_{A}+e_{H}-\delta_{H}e_{H} \right)^{3}d_{A}u_{H}+f^{2}(-\left( -1+\delta_{A} \right)\left( 6+\left( -1+\delta_{H} \right)e_{H}\left( 11+\delta_{A}+5\left( -1+\delta_{H} \right)e_{H} \right) \right)d_{H}u_{A}+e_{H}\left( 1-\delta_{A}+\left( -1+\delta_{H} \right)e_{H} \right)\left( -\left( -1+\delta_{A} \right)^{2}+\left( -1+\delta_{H} \right)^{2}e_{H} \right)d_{A}u_{H})+(-1+\delta_{A}+e_{H}-\delta_{H}e_{H})f(\left( 6+\left( -1+\delta_{H} \right)e_{H}\left( 6+\left( -1+\delta_{H} \right)e_{H} \right) \right)d_{H}u_{A}-2\left( -1+\delta_{A} \right)e_{H}\left( -1+\delta_{A}+e_{H}-\delta_{H}e_{H} \right)d_{A}u_{H})))$$

$$k_{11}=-2{a_{A}}^{2}{a_{H}}^{3}E_{1}{e_{H}}^{3}fd_{A}{d_{H}}^{2}\left( a_{H}e_{H}\left( 2+\left( -1+\delta_{H} \right)e_{H}+\left( -1+\delta_{H} \right)f^{2} \right)I+\left( \left( 1+\left( -1+\delta_{H} \right)e_{H} \right)\left( 2+\left( -1+\delta_{H} \right)e_{H}+f\left( -2+f-\delta_{H}f \right) \right)+\delta_{A}\left( 2\left( -1+f \right)-\left( -1+\delta_{H} \right)\left( e_{H}-2e_{Hf}+f^{2} \right) \right) \right)d_{A}d_{H} \right)u_{A}$$

$$k_{12}=-a_{A}{a_{H}}^{4}E_{1}{e_{H}}^{4}fd_{A}d_{H}\left( a_{H}e_{H}I+\left( 1+\left( -1+\delta_{H} \right)e_{H}+\delta_{A}\left( -1+f \right)+f\left( -1+f-\delta_{H}f \right) \right)d_{A}d_{H} \right)u_{A}$$

$$m_{6}=a_{H}e_{H}$$

$$m_{7}= a_{A}\left( 1+\left( -1+\delta_{H} \right)e_{H} \right)d_{H}$$

$$m_{8}=a_{A}(a_{H}e_{H}I-\left( -1+\delta_{A}+e_{H}-\delta_{H}e_{H} \right)d_{A}d_{H})$$

$$m_{9}= d_{A}+\left( -1+\delta_{A} \right)fd_{A}$$

$$m_{10}= (1+(-1+\delta_{H})f^{2})$$

$$m_{11}= a_{A}a_{H}e_{H}I-a_{A}\delta_{A}d_{A}d_{H}$$

$$m_{12}=a_{H}e_{H}$$

$$m_{13}=a_{A}\left( 1+\left( -1+\delta_{H} \right)e_{H} \right)d_{H}$$

$$m_{14}=a_{A}a_{H}e_{H}(a_{H}e_{H}I-\left( -1+\delta_{A}+e_{H}-\delta_{H}e_{H} \right)d_{A}d_{H})$$

$$m_{15}=d_{A}+\left( -1+ \delta_{A} \right)fd_{A}$$

$$m_{16}=1+\left( -1+ \delta_{H} \right)f^{2}$$

$$m_{17}=a_{A}a_{H}e_{H}I-a_{A}\delta_{A}d_{A}d_{H}$$

**Contaminant Concentrations:**

$$y_{1}=e_{a}d_{A}d_{H}(\left( a_{H}e_{H}l+a_{A}\left( 1+\left( -1+\delta_{H} \right)e_{H} \right)d_{H} \right)\left( -a_{H}e_{H}\left( -1+\delta_{A}+f-\delta_{H}f \right)l+a_{A}\left( -\left( -1+\delta_{A} \right)\left( 1+\left( -1+\delta_{H} \right)e_{H} \right)+d_{A}\left( -1+\delta_{H} \right)f \right)d_{H} \right)u_{A}+a_{A}\left( -1+\delta_{H} \right)e_{H}\left( 1-\delta_{A}+\left( -1+\delta_{H} \right)e_{H} \right)d_{A}\left( a_{H}e_{H}l+a_{A}\delta_{A}d_{H} \right)u_{H}$$

$$y_{2}=a_{A}a_{H}e_{H}(d_{H}\left( a_{H}e_{H}l+a_{A}\left( 1+\left( -1+\delta_{H} \right)e_{H} \right)d_{H} \right)\left( -E_{0}\left( -1+\delta_{A}+e_{H}-\delta_{H}e_{H} \right)d_{A}-\left( -1+\delta_{H} \right)E_{1}f u_{A} \right)+\left( -1+\delta_{H} \right)E_{1}e_{H}d_{A}\left( a_{H}e_{H}l+a_{A}\left( -1+2\delta_{A}+e_{H}-\delta_{H}e_{H} \right)d_{H} \right)u_{H}$$

$$y_{3}=a_{A}a_{H}e_{H}d_{A}d_{H}(a_{H}e_{H}\left( 2+2\left( -1+\delta_{H} \right)e_{H}+\delta_{A}\left( -2+f \right)+f\left( -1+f-\delta_{H}f \right) \right)l+a_{A}(\delta_{A}\left( -2+f+\left( -1+\delta_{H} \right)e_{H}f-2\left( -1+\delta_{H}f^{2} \right)+\left( 1+\left( -1+\delta_{H} \right)e_{H} \right)\left( 2+f\left( -1+\left( -1+\delta_{H} \right)f \right) \right) \right)d_{H}$$

$$y_{4}=E_{1}\left( a_{H}e_{H}l+a_{A}d_{H} \right)\left( a_{H}e_{H}l+a_{A}\left( 1+\left( -1+\delta_{H} \right)e_{H} \right)d_{H} \right)u_{A}+a_{A}\left( -1+\delta_{H} \right)E_{1}e_{H}fd_{A}\left( a_{H}e_{H}l+a_{A}\delta_{A}d_{H} \right)u_{H}$$

$$y_{5}=a_{A}a_{H}e_{H}f(a_{H}E_{0}e_{H}l+a_{A}E_{0}\left( 1+\left( -1+\delta_{H} \right)e_{H} \right)d_{H}-a_{A}\left( -1+\delta_{H} \right)E_{1}e_{H}u_{H}$$

$$y_{6}=E_{1}d_{H}(a_{H}^{2}e_{H}^{2}f l^{2}u_{A}+a_{A}a_{H}e_{H}l\left( \left( 2+\left( -1+\delta_{H} \right)e_{H} \right)fd_{H}u_{A}+e_{H}\left( 1+\left( -1+\delta_{H} \right)e_{H}+\delta_{A}\left( -1+f \right)-f \right)d_{A}u_{H}+a_{A}^{2}d_{H}\left( e_{H}\left( 1-\delta_{A}+\left( -1+\delta_{H} \right)e_{H} \right)d_{A}u_{H}+\left( 1+\left( -1+\delta_{H} \right)e_{H} \right)f\left( d_{H}u_{A}+\left( -1+\delta_{A} \right)e_{H}d_{A}u_{H} \right) \right) \right)$$

$$y_{7}=a_{A}a_{H}e_{H}(a_{H}e_{H}l\left( E_{0}f^{2}d_{H}+E_{1}e_{H}u_{H} \right)+a_{A}d_{H}(E_{0}\left( 1+\left( -1+\delta_{H} \right)e_{H} \right)f^{2}d_{H}+E_{1}e_{H}u_{H}$$

$$x_{1}= -a_{A}\left( -1+\delta_{A}+e_{H}-\delta_{H}e_{H} \right)d_{A}^{2}d_{H}^{2}(a_{H}e_{h}\left( 1+\left( -1+\delta_{H} \right)e_{H}+\delta_{A}\left( -1+f \right)+f\left( -1+f-\delta_{H}f \right) \right)l+a_{A}\left( -\left( 1+\left( -1+\delta_{H} \right)e_{H} \right)\left( -1+f \right)+\delta_{A}\left( -1+f\left( 1+\left( -1+\delta_{H} \right)e_{H}+f-\delta_{H}f \right) \right) \right)d_{H}$$

$$x_{2}=a_{A}a_{H}e_{H}d_{A}d_{H}(a_{H}e_{H}\left( 2+2\left( -1+\delta_{H} \right)e_{H}+\delta_{A}\left( -2+f \right)+f\left( -1+f-\delta_{H}f \right) \right)l+a_{A}\left( \delta_{A}\left( -2+f+\left( -1+\delta_{H} \right)e_{H}f-2\left( -1+\delta_{H} \right)f^{2} \right)+\left( 1+\left( -1+\delta_{H} \right)e_{H} \right)\left( 2+f\left( -1+\left( -1+\delta_{H} \right)f \right) \right) \right)d_{H}$$

$$x_{3}=a_{A}a_{H}^{2}e_{H}^{2}(a_{H}e_{H}l+a_{A}\left( 1+\left( -1+\delta_{H} \right)f^{2} \right)d_{H})$$

$$x_{4}=a_{A}d_{A}d_{H}(a_{H}e_{H}\left( 1+\left( -1+\delta_{H} \right)e_{H}+\delta_{A}\left( -1+f \right)+f\left( -1+f+\delta_{H}f \right) \right)l+a_{A}\left( -\left( 1+\left( -1+\delta_{H} \right)e_{H} \right)\left( -1+f \right)+\delta_{A}\left( -1+f\left( 1+\left( -1+\delta_{H} \right)e_{H}+f-\delta_{H}f \right) \right) \right)d_{H}$$

$$x_{5}=a_{A}a_{H}e_{H}(a_{H}e_{H}l+a_{A}\left( 1+\left( -1+\delta_{H} \right)f^{2} \right)d_{H})$$

$$x_{6}=a_{A}e_{H}d_{A}d_{H}^{2}(a_{H}e_{H}\left( 1+\left( -1+\delta_{H} \right)e_{H}+\delta_{A}\left( -1+f \right)+f\left( -1+f-\delta_{H}f \right) \right)l+a_{A}\left( -\left( 1+\left( -1+\delta_{H} \right)e_{H} \right)\left( -1+f \right)+\delta_{A}\left( -1+f\left( 1+\left( -1+\delta_{H} \right)e_{H}+f-\delta_{H}f \right) \right) \right)d_{H}$$

$$x_{7}=a_{A}a_{H}e_{H}^{2}d_{H}(a_{H}e_{H}l+a_{A}\left( 1+\left( -1+\delta_{H} \right)f^{2} \right)d_{H})$$

$$v_{1}=a_{A}^{2}(a_{H}^{2}e_{H}^{2}I^{2}\left( E_{0}\left( 1+\left( -1+\delta_{H} \right)e_{H} \right)d_{H}-\left( -1+\delta_{H} \right)E_{1}e_{H}u_{H} \right)+a_{H}e_{H}Id_{H}(\left( 1+\left( -1+\delta_{H} \right)e_{H} \right)d_{H}\left( -E_{0}\left( -1+\delta_{A}+e_{H}-\delta_{H}e_{H} \right)d_{A}-\left( -1+\delta_{H} \right)E_{1}fu_{A} \right)-\left( -1+\delta_{H} \right)E_{1}e_{H}\left( 1-2\delta_{A}+\left( -1+\delta_{H} \right)e_{H} \right)d_{A}u_{H}+E_{1}d_{A}d_{H}^{2}\left( -\left( 1+\left( -1+\delta_{H} \right)e_{H} \right)\left( \left( -1+\delta_{A} \right)\left( 1+\left( -1+\delta_{H} \right)e_{H} \right)-\delta_{A}\left( -1+\delta_{H} \right)f \right)d_{H}u_{A}-\delta_{A}\left( -1+\delta_{H} \right)e_{H}\left( -1+\delta_{A}+e_{H}-\delta_{H}e_{H} \right)d_{A}u_{H} \right))$$

$$v_{2}=a_{A}a_{H}e_{H}(a_{H}^{2}E_{0}e_{H}^{2}I^{2}+E_{1}d_{H}d_{A}\left( \left( -2\left( -1+\delta_{A} \right)\left( 1+\left( -1+\delta_{H} \right)e_{H} \right)+\left( -1+\delta_{H} \right)\left( 1+\delta_{A}+\left( -1+\delta_{H} \right)e_{H} \right)f \right)d_{H}u_{A}-\left( -1+\delta_{H} \right)e_{H}\left( -1+\delta_{A}+e_{H}-\delta_{H}e_{H} \right)d_{A}u_{H} \right)+a_{H}e_{H}I\left( -E_{0}\left( -1+\delta_{A}+e_{H}-\delta_{H}e_{H} \right)d_{A}d_{H}-\left( -1+\delta_{H} \right)E_{1}\left( fd_{H}u_{A}-e_{H}d_{A}u_{H} \right) \right))$$

$$v_{3}= -a_{H}^{2} E_{1}e_{H}^{2}\left( -1+\delta_{A}+f-\delta_{H}f \right)d_{A}d_{H}u_{A}$$

$$v_{4}=a_{H}^{2}e_{1}e_{H}^{2}l^{2}u_{A}+a_{A}^{2}(\left( 1+\left( -1+\delta_{H} \right)e_{H} \right)d_{H}\left( a_{H}E_{0}e_{H}fI+E_{1}d_{H}u_{A} \right)-\left( -1+\delta_{H} \right)E_{1}e_{H}f\left( a_{H}e_{H}I-\delta_{A}d_{A}d_{H} \right)u_{H}$$

$$v_{5}=a_{A}a_{H}e_{H}(a_{H}E_{0}+e_{H}fI+2E_{1}d_{H}u_{A}+\left( -1+\delta_{H} \right)E_{1}e_{H}\left( d_{H}u_{A}+fd_{A}u_{H} \right))$$

$$v_{6}=d_{H}(a_{H}^{2}E_{1}e_{H}^{2}fl^{2}u_{A}+a_{A}^{2}\left( \left( 1+\left( -1+\delta_{H} \right)e_{H} \right)fd_{H}\left( a_{H}E_{0}e_{H}fI+E_{1}d_{H}u_{A} \right)+E_{1}e_{H}\left( a_{H}e_{H}I+\left( 1-\delta_{A}-e_{H}+\delta_{H}e_{H}+\left( -1+\delta_{A} \right)\left( 1+\left( -1+\delta_{H} \right)e_{H} \right)f \right)d_{A}d_{H} \right)u_{H} \right))$$

$$v_{7}=a_{A}a_{H}e_{H}(a_{H}e_{H}I\left( E_{0}f^{2}d_{H}+E_{1}e_{H}u_{H} \right)+E_{1}d_{H}(\left( 2+\left( -1+\delta_{H} \right)e_{H} \right)fd_{H}u_{A}+e_{H}\left( 1+\left( -1+\delta_{H} \right)e_{H}+\delta_{A}\left( -1+f \right)-f \right)d_{A}u_{H}$$

$$b_{1}=a_{A}^{2}d_{H}\left( a_{H}e_{H}I-\left( -1+\delta_{A}+e_{H}-\delta_{H}e_{H} \right)d_{A}d_{H} \right)\left( a_{H}e_{H}\left( 1+\left( -1+\delta_{H} \right)f^{2} \right)I+\left( -\left( 1+\delta_{H} \right)e_{H} \right)\left( -1+f \right)+\delta_{A}\left( -1+f\left( 1+\left( -1+\delta_{H} \right)e_{H}+f-\delta_{H}f \right) \right) \right)d_{A}d_{H}$$

$$b_{2}=a_{A}a_{H}e_{H}(a_{H}e_{H}I-\left( -1+\delta_{A}+e_{H}-\delta_{H}e_{H} \right)d_{A}d_{H})(a_{H}e_{H}I+\left( 1+\left( -1+\delta_{H} \right)e_{H}+\delta_{A}\left( -1+f \right)+f\left( -1+f-\delta_{H}f \right) \right)d_{A}d_{H})$$

$$b_{3}=a_{A}^{2}d_{H}(a_{H}e_{H}\left( 1+\left( -1+\delta_{H} \right)f^{2} \right)I-\left( 1+\left( -1+\delta_{H} \right)e_{H} \right)\left( -1+f \right)d_{A}d_{H}+\delta_{A}d_{A}\left( -\delta_{H}f^{2}+\left( -1+f\left( 1+\delta_{H} \right)e_{H}+f \right) \right)d_{H}))$$

$$b_{4}=a_{A}a_{H}e_{H}(a_{H}e_{H}I+\left( 1+\left( -1+\delta_{H} \right)e_{H}+\delta_{A}\left( -1+f \right)+f\left( -1+f-\delta_{H}f \right) \right)d_{A}d_{H}))$$

$$b_{5}=a_{A}^{2}e_{H}d_{H}^{2}(a_{H}e_{H}\left( 1+\left( -1+\delta_{H} \right)f^{2} \right)I+\left( -\left( 1+\left( -1+\delta_{H} \right)e_{H} \right)\left( -1+f \right)+\delta_{A}\left( -1+f\left( 1+\left( -1+\delta_{H} \right)e_{H}+f-\delta_{H}f \right) \right) \right)d_{A}d_{H}$$

$$b_{6}=a_{A}a_{H}e_{H}^{2}d_{H}(a_{H}e_{H}I+\left( 1+\left( -1+\delta_{H} \right)e_{H}+\delta_{A}\left( -1+f \right)+f\left( -1+f-\delta_{H}f \right) \right)d_{A}d_{H})$$
