## Supplemental Information 4 for "Integrating ecosystem and contaminant models to predict the effects of ecosystem fluxes on contaminant dynamics"

**Appendix S4: Additional Figures and Table**

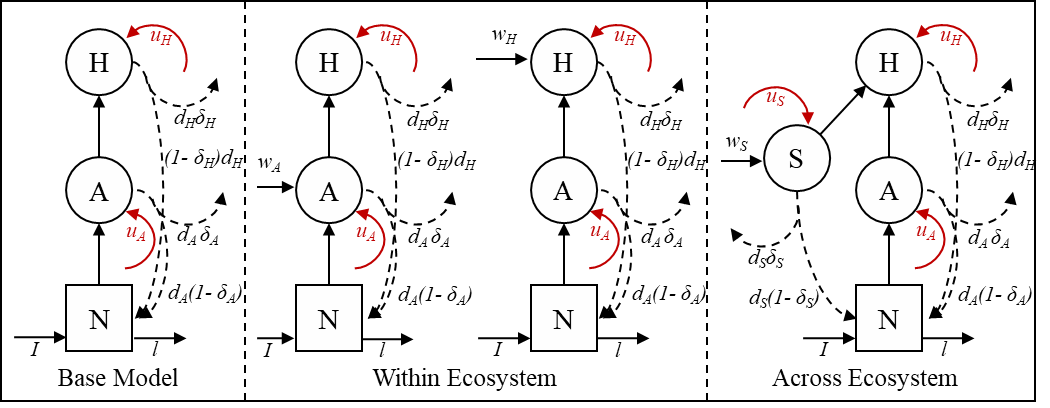

**Figure S1.** Diagram of ecosystem models analyzed here where *N* represents inorganic nutrient stocks, *A* autotroph stocks within ecosystems, *H* herbivore stocks within ecosystems, and *S* autotroph subsidies across ecosystems (i.e., movement from a different ecosystem); *I* indicates the input rate of inorganic nutrients, *l* indicates the loss rate of inorganic nutrients, *w*_i_ indicates input rate of trophic level *i* (where *i* is *N*, *A*, or *H*), *u_i_* indicates the environmental uptake of contaminant by trophic level *i*, and *d_i_* indicates the proportion of biomass lost from trophic level *i* and 1– *δ_i_* is the portion of this loss that is recycled. See model description in text for full model details. Circles and squares reflect biomass and contaminant mass in each trophic level, while black lines indicate parallel flows of nutrients and contaminants between compartments, dashed black lines indicate the parallel nutrients and contaminants lost and recycled back to the inorganic nutrient pool, and red lines indicate flows relevant to contaminants only (i.e., red line flows are only included in contaminant model). The base model considers only biomass transfer along the food chain in a single patch, open at the basal level for flows of inorganic nutrients in (*I*) and out (*l*). The within ecosystem model considers fluxes of biomass from donor ecosystems of the same type – for example the movement of phytoplankton between two pond ecosystems. The across ecosystem model considers fluxes of biomass from donor ecosystems of a different type – for example litter fall from terrestrial systems into aquatic systems or marine wrack on beaches.

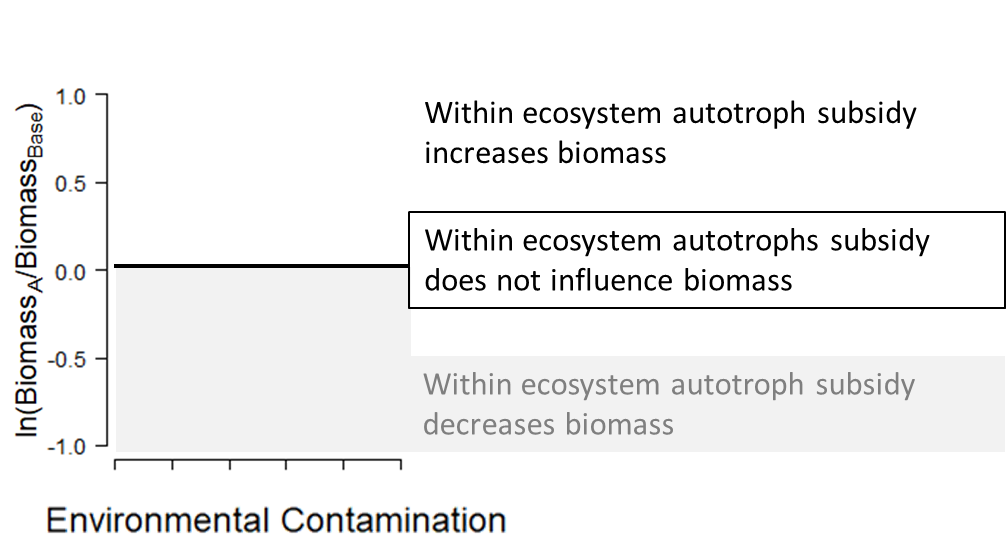

**Figure S2.** A conceptual diagram for how to interpret the natural log response variables. In this example the dynamics of within ecosystem autotroph subsidies compared to the base model (no-movement scenario) are being compared. A value greater than zero indicates that within ecosystem autotroph subsidies increases biomass of chosen trophic level as compared to no-movement dynamics, while a value less than zero indicates that movement of autotrophs within ecosystems decreases biomass as compared to a no-movement model.

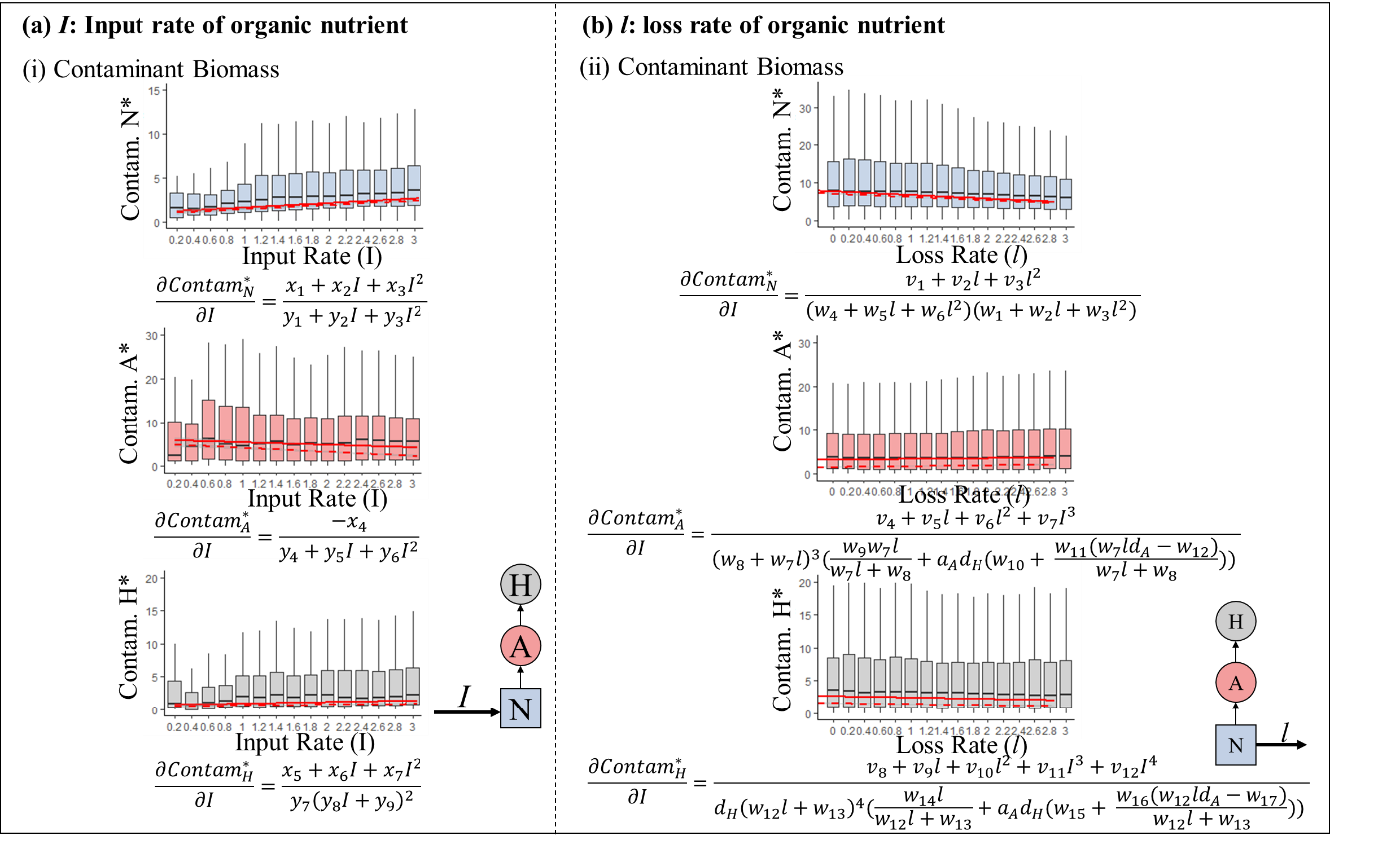

**Figure S3.** Demonstration of how (a) input rate of organic nutrients (*I*) and (*b*) loss rate of organic nutrients (*l*) influence equilibrium contaminant mass with the range of parameters described in the main text. While these results are dependent on the parameters selected, the simplified partial derivatives below each graph demonstrate that input rate and loss rate influence equilibrium contaminant concentrations irrespective of parameter values. Overlaid on (i & ii) are two representative chemicals – polychlorinated biphenyls (solid line in red) and cyclic methyl siloxanes (dashed line in red) for the same parameter values but have chemical specific values of environmental uptake of contaminant by trophic level *i* (*u_i_*) and assimilation efficiencies of the contaminant in biotic compartment (*f*). For more details on the substitutions for the *c_i_*s, *g_i_*s, *k_i_*s, and *m_i_*s see Appendix 2. For full equilibria solutions see Appendix 2.

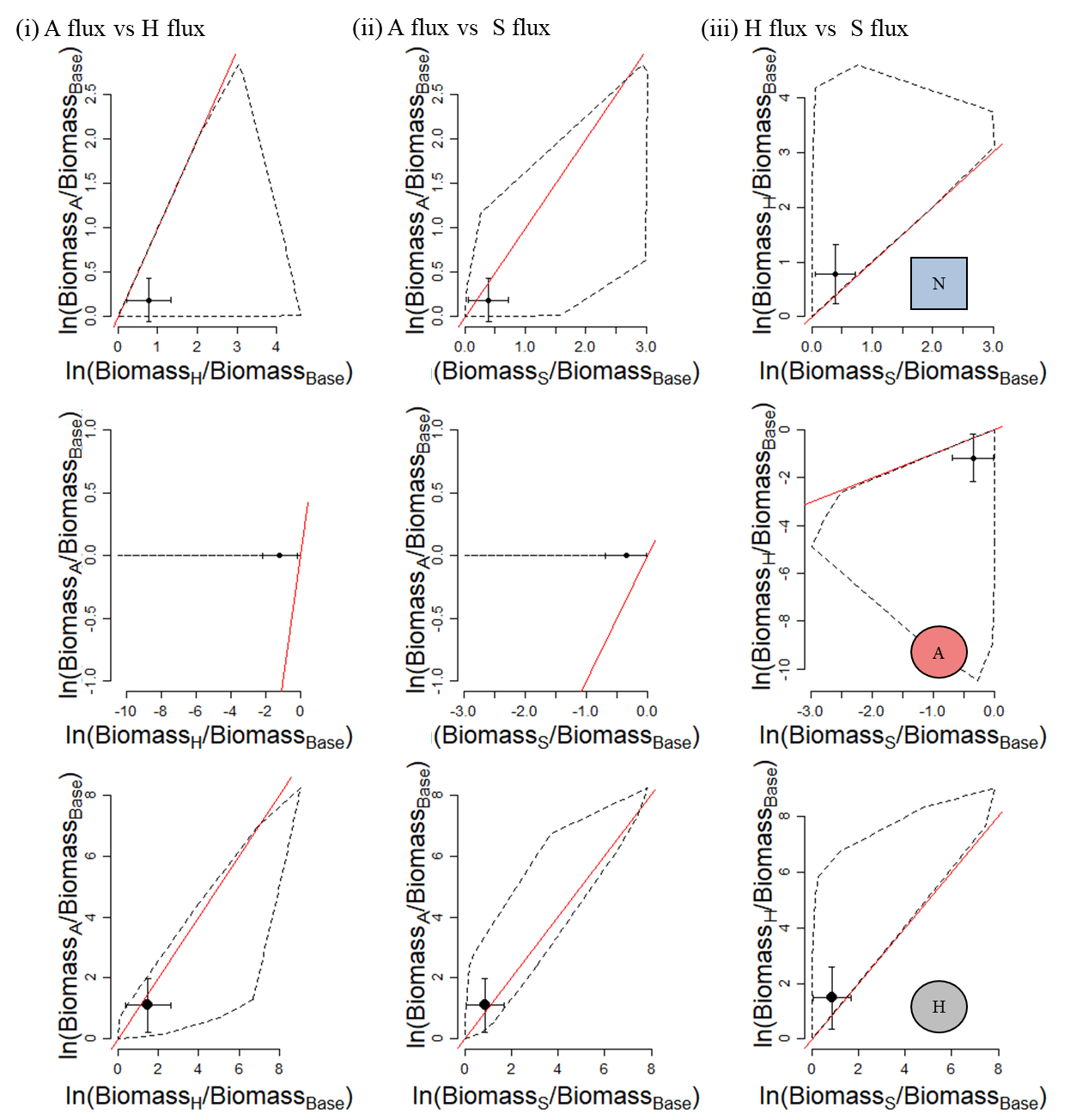

**Figure S4**. Attenuation plots comparing the influence of (i) within ecosystem autotroph fluxes on biomass to within ecosystem herbivore subsidies, (ii) within ecosystem autotroph influx to across ecosystem autotroph fluxes and (iii) within ecosystem herbivore influx to across ecosystem autotroph fluxes. The red line indicates a one to one relationship and everything above and to the left of the red line means the y-axis has a stronger impact than the x-axis, while everything below and to the right of the red line means the x-axis has a stronger impact than the y-axis (see Figure 5 for figure for contaminant concentration, and Appendix 4: Figure S7 for contaminant mass). The dots are the mean value from simulations with standard deviation confidence bars and dashed lines representing 95% confidence hulls.

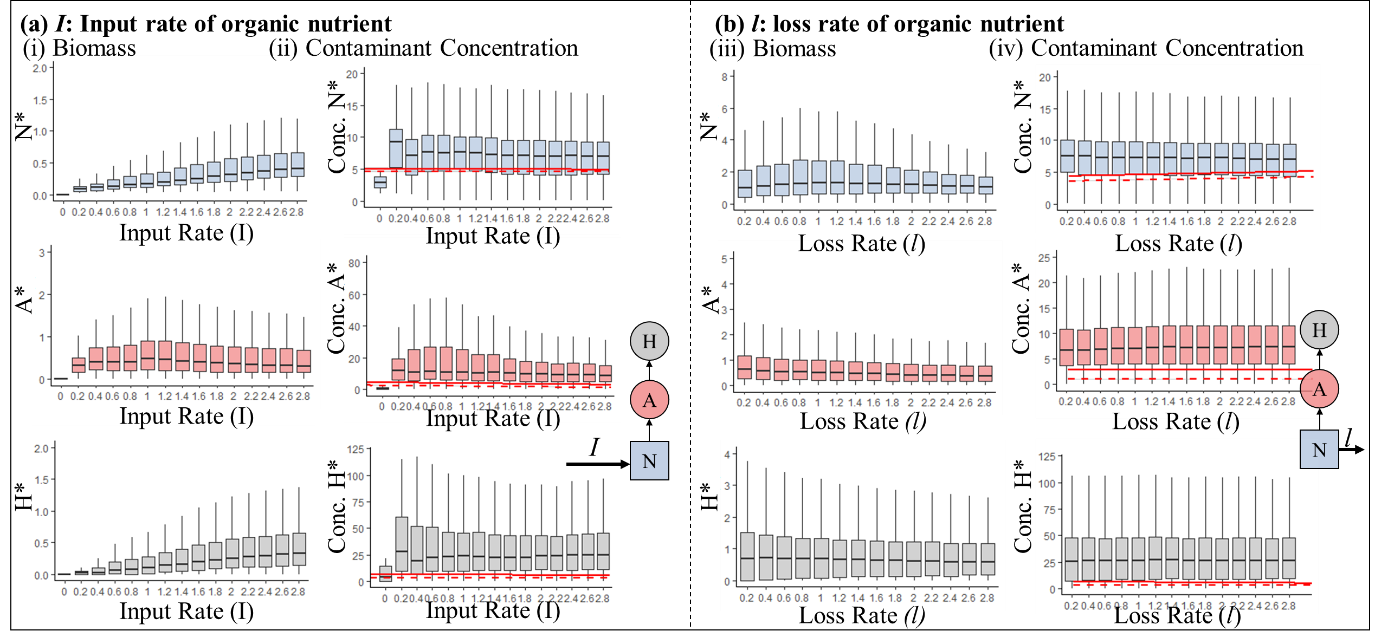

**Figure S5**. Demonstration of how (a) input rate of organic nutrients (*I*) and (b) loss rate of organic nutrients (*l*) influence the (i & iii) equilibrium biomass and (ii & iv) equilibrium contaminant concentration for $\alpha_{1}$ = 1 and *LC_50_* = 80 with the range of parameters described in the main text and the modified mortality rate described in Appendix S6. Overlaid on (ii & iv) are two representative chemicals – polychlorinated biphenyls (solid line in red) and cyclic methyl siloxanes (dashed line in red) for the same parameter values, but have chemical specific values of environmental uptake of contaminant by trophic level *i* (*u_i_*) and assimilation efficiencies of the contaminant in biotic compartment (*f*). This figure is directly comparable to Figure 2.

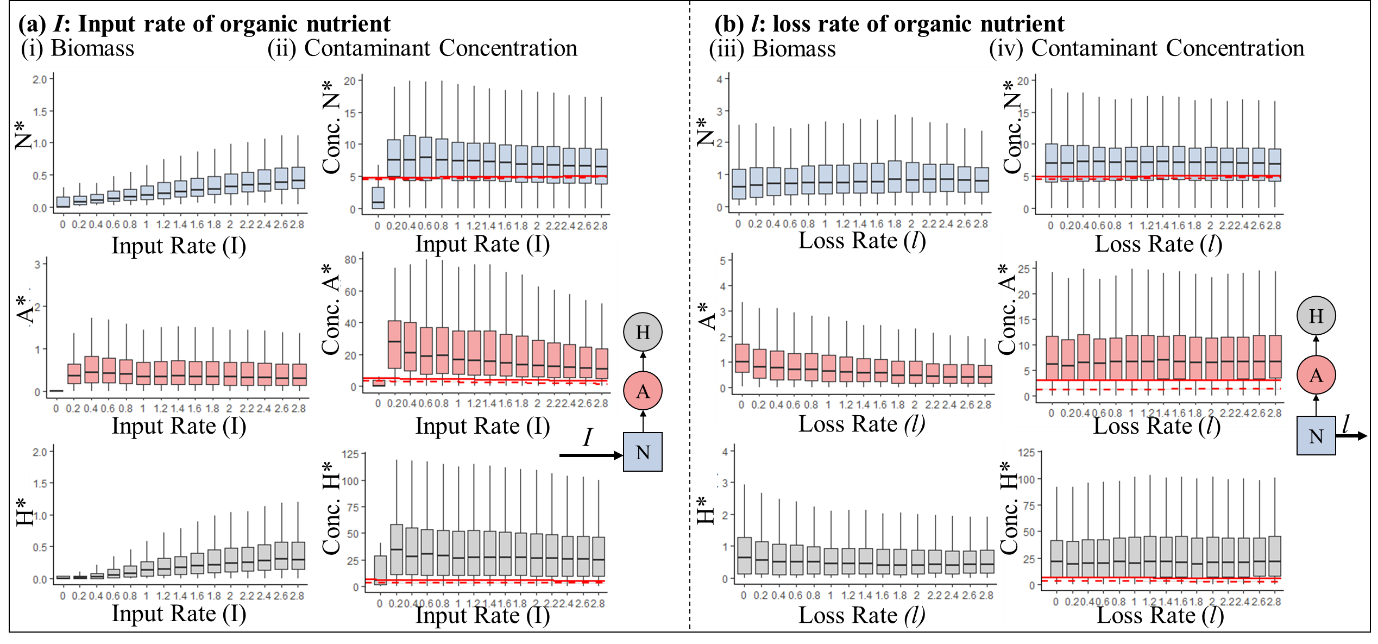

**Figure S6.** Demonstration of how (a) input rate of organic nutrients (*I*) and (b) loss rate of organic nutrients (*l*) influence the (i & ii) equilibrium biomass and (iii & iv) equilibrium contaminant concentration for $\alpha_{1}$ = 4 and *LC_50_* = 80 with the range of parameters described in the main text and the modified mortality rate described in Appendix S6. Overlaid on (ii & iv) are two representative chemicals – polychlorinated biphenyls (solid line in red) and cyclic methyl siloxanes (dashed line in red) for the same parameter values, but have chemical specific values of environmental uptake of contaminant by trophic level *i* (*u_i_*) and assimilation efficiencies of the contaminant in biotic compartment (*f*). This figure is directly comparable to Figure 2.

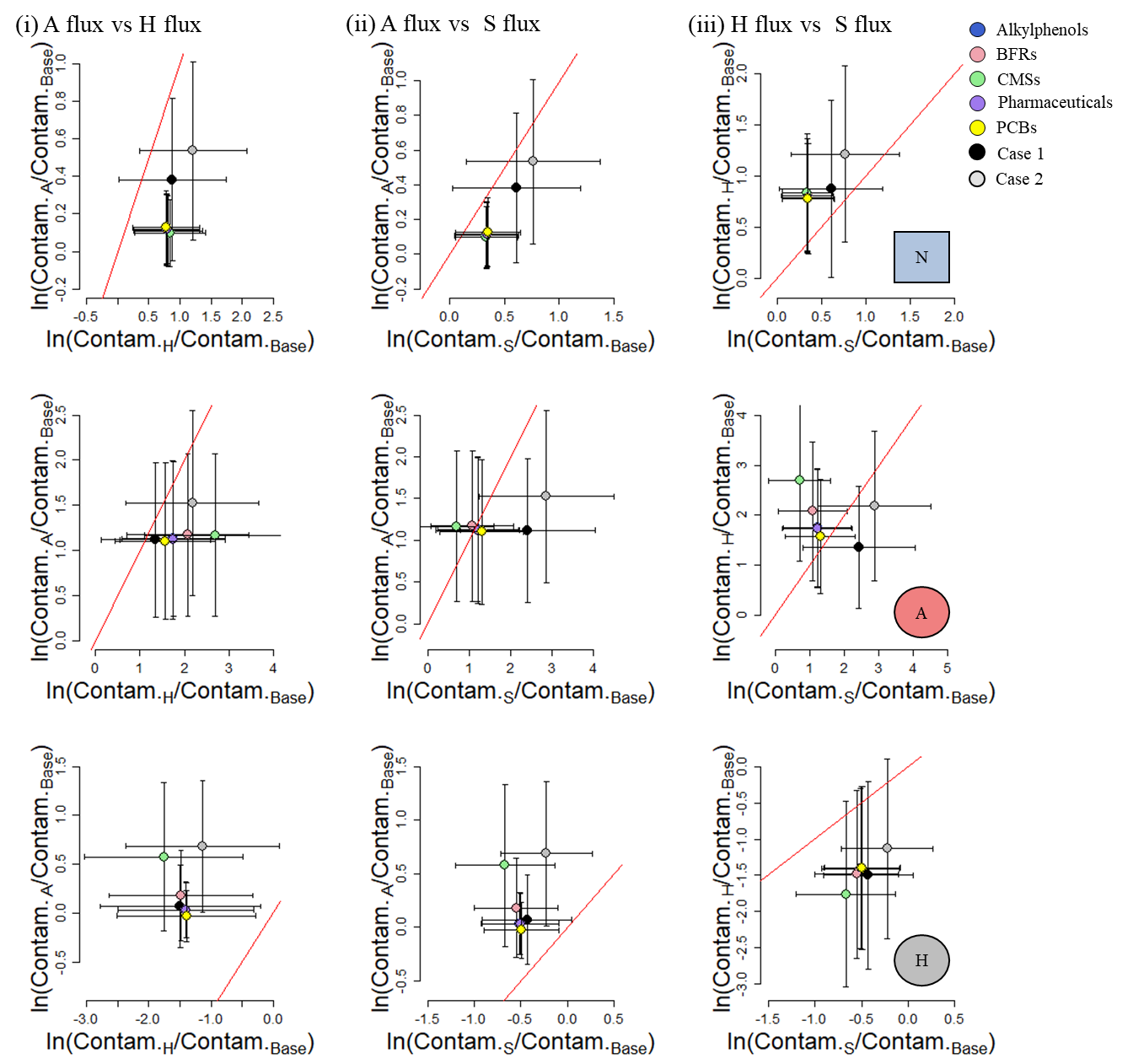

**Figure S7.** Attenuation plots comparing the influence of (i) within ecosystem autotroph fluxes on contaminant mass to within ecosystem herbivore fluxes, (ii) within ecosystem autotroph fluxes to across ecosystem autotroph fluxes and (iii) within ecosystem herbivore influx to across ecosystem autotroph fluxes. Case 1 is when the recipient ecosystem is more contaminated than the donor ecosystem for both the x and y-axes, and Case 2 is when the recipient ecosystem is less contaminated than the donor ecosystem. The red line indicates a one to one relationship and everything above and to the left of the red line means the y-axis has a stronger impact than the x-axis, while everything below and to the right of the red line means the x-axis has a stronger impact than the y-axis (see Appendix 4: Figure S4 for plot of biomass and Appendix 4: Figure S8 for plot of contaminant concentration). The dots are the mean value from simulations with standard deviation confidence bars. Coloured dots are the mean values from simulations when specific chemical properties are used in the simulations (for ranges of values see Appendix 4: Table S1).

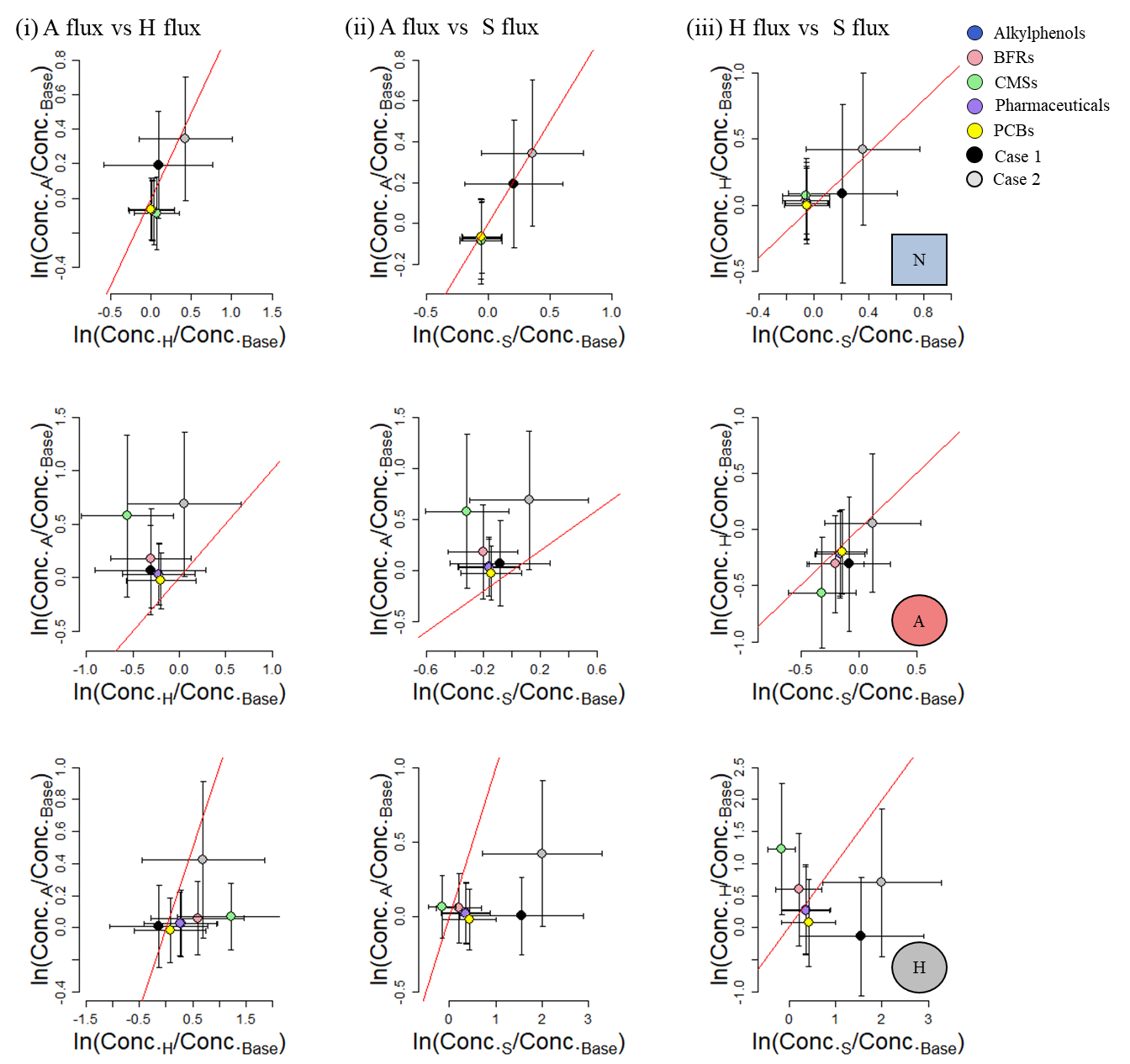

**Figure S8.** Attenuation plots comparing the influence of (i) within ecosystem autotroph fluxes on contaminant concentration to within ecosystem herbivore fluxes, (ii) within ecosystem autotroph fluxes to across ecosystem autotroph fluxes and (iii) within ecosystem herbivore influx to across ecosystem autotroph fluxes. Case 1 is when the recipient ecosystem is more contaminated than the donor ecosystem, and Case 2 is when the recipient ecosystem is less contaminated than the donor ecosystem. The red line indicates a one to one relationship and everything above and to the left of the red line means the y-axis has a stronger impact than the x-axis, while everything below and to the right of the red line means the x-axis has a stronger impact than the y-axis. The dots are the mean value from simulations with standard deviation confidence bars. Coloured dots are the mean values from simulations when specific chemical properties are used in the simulations (for ranges of values see Appendix 4: Table S1).

**Table S1.** Ranges of chemical properties used in simulations for each class of chemical plotted on Figure S7 and S8 in Appendix 4. Chemical properties were taken from Walters et al. (2016) and converted to environmental uptakes (*u_i_*) and assimilation efficiencies (*f*) using the equations provided in Arnot and Gobas (2004). Note: there is not much variability in environmental uptake parameters because the equation used to calculate this parameter is relatively invariant to logK_OW_.

|  | *f* | | *u_i_* | |
| --- | --- | --- | --- | --- |
|  | min | max | min | max |
| Alkylphenols | 0.268 | 0.339 | 0.540 | 0.540 |
| Brominated Flame Retardants | 0.004 | 0.404 | 0.540 | 0.541 |
| Cyclic Methyl Siloxanes | 0.003 | 0.049 | 0.541 | 0.541 |
| Pharmaceuticals and Personal Care Products | 0.257 | 0.356 | 0.540 | 0.540 |
| Polychlorinated Biphenyls | 0.002 | 0.479 | 0.539 | 0.541 |
