## Supplemental Information 5 for "Integrating ecosystem and contaminant models to predict the effects of ecosystem fluxes on contaminant dynamics"

**Appendix S5: Demonstration of how donor ecosystem contaminant concentrations impact equilibrium concentrations**

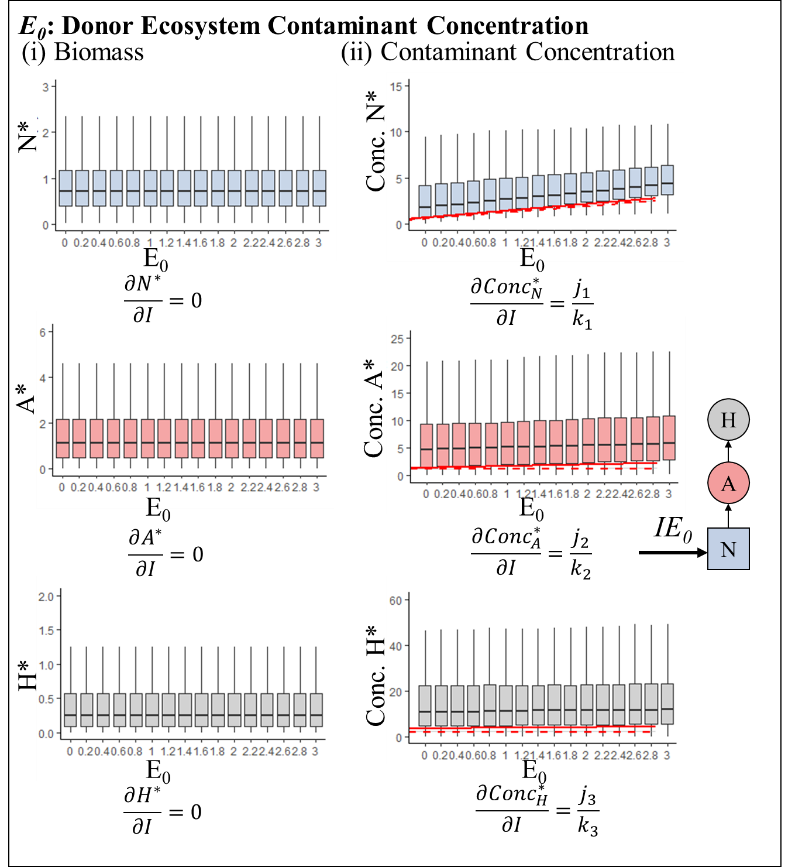

**Figure S1.** Demonstration of how donor ecosystem contaminant concentration (*E_0_*) influences the (i) equilibrium biomass and (ii) equilibrium contaminant concentration with the range of parameters described in the main text. While these results are dependent on the parameters selected, the simplified partial derivatives below each graph demonstrate that donor ecosystem contaminant concentration has a linear impact on equilibrium contaminant concentrations irrespective of parameter values. Overlaid on (ii) are two representative chemicals – polychlorinated biphenyls (solid line in red) and cyclic methyl siloxanes (dashed line in red) for the same parameter values but have chemical specific values of environmental uptake of contaminant by trophic level *i* (*u_i_*) and assimilation efficiencies of the contaminant in biotic compartment (*f*). For more details on the substitutions for the *j_i_*s and *k_i_*s see below substitutions. For full equilibria solutions see Appendix 1.

**Substitutions for the partial derivatives presented in Fig. 1.**

**Contaminant Biomass:**

$$j_{1}=a_{H}e_{H}I (a_{H}e_{H}I-\left( -1+\delta_{A}+e_{H}-\delta_{H}e_{H} \right)d_{A}d_{H}))/(a_{H}^{2}e_{H}^{2}Il+a_{H}e_{H}\left( a_{A}\left( 1+\left( -1+\delta_{H} \right)f^{2} \right)I-\left( -1+\delta_{A}+e_{H}-\delta_{H}e_{H}+f-\delta_{A}f+\left( -1+ \delta_{H} \right)f^{2} \right)ld_{A} \right)d_{H}+a_{A}\left( -\left( \left( 1+\left( -1+\delta_{H} \right)e_{H} \right)\left( -1+f \right) \right)+ \delta_{A}\left( -1+f\left( 1+\left( -1+\delta_{H} \right)e_{H}+f-\delta_{H}f \right) \right) \right)d_{A}d_{H}^{2})$$

$$j_{2}=fId_{H}(a_{H}e_{H}l+a_{A}\left( 1+\left( -1+\delta_{H} \right)e_{H} \right)d_{H})/(a_{H}^{2}e_{H}^{2}I l+a_{H}e_{H}\left( a_{A}\left( 1+\left( -1+\delta_{H} \right)f^{2} \right)I-\left( -1+\delta_{A}+e_{H}-\delta_{H}e_{H}+f-\delta_{A}f+\left( -1+\delta_{H} \right)f^{2} \right)l d_{A} \right)d_{H}+a_{A}(-\left( \left( 1+\left( -1+\delta_{H} \right)e_{H}\left( -1+f \right) \right)+\delta_{A}\left( -1+f\left( 1+\left( -1+\delta_{H} \right)e_{H}+f-\delta_{H}f \right) \right) \right)d_{A}d_{H}^{2})$$

$$j_{3}=(f^{2}I \left( a_{A}a_{H}e_{H}I-a_{H}e_{H}ld_{A}-a_{A}\delta_{A}d_{A}d_{H} \right))/(a_{H}^{2}e_{H}^{2}I l+a_{H}e_{H}\left( a_{A}\left( 1+\left( -1+\delta_{H} \right)f^{2} \right)I-\left( -1+\delta_{A}+e_{H}-\delta_{H}e_{H}+f-\delta_{A}f+\left( -1+\delta_{H} \right)f^{2} \right)l d_{A} \right)d_{H}+a_{A}(-\left( \left( 1+\left( -1+\delta_{H} \right)e_{H}\left( -1+f \right) \right)+ \delta_{A}\left( -1+f\left( 1+\left( -1+\delta_{H} \right)e_{H}+f-\delta_{H}f \right) \right) \right)d_{A}d_{H}^{2}$$

$$k_{1}=a_{H}e_{H}I (a_{H}e_{H}l+a_{A}\left( 1+(-1+\delta_{H})e_{H} \right)d_{H}))/(a_{H}^{2}e_{H}^{2}Il+a_{H}e_{H}\left( a_{A}\left( 1+\left( -1+\delta_{H} \right)f^{2} \right)I-\left( -1+\delta_{A}+e_{H}-\delta_{H}e_{H}+f-\delta_{A}f+\left( -1+ \delta_{H} \right)f^{2} \right)ld_{A} \right)d_{H}+a_{A}\left( -\left( \left( 1+\left( -1+\delta_{H} \right)e_{H} \right)\left( -1+f \right) \right)+ \delta_{A}\left( -1+f\left( 1+\left( -1+\delta_{H} \right)e_{H}+f-\delta_{H}f \right) \right) \right)d_{A}d_{H}^{2})$$

$$k_{2}=a_{H}e_{H}fI (a_{H}e_{H}l+a_{A}\left( 1+(-1+\delta_{H})e_{H} \right)d_{H}))/(a_{H}^{2}e_{H}^{2}Il+a_{H}e_{H}\left( a_{A}\left( 1+\left( -1+\delta_{H} \right)f^{2} \right)I-\left( -1+\delta_{A}+e_{H}-\delta_{H}e_{H}+f-\delta_{A}f+\left( -1+ \delta_{H} \right)f^{2} \right)ld_{A} \right)d_{H}+a_{A}\left( -\left( \left( 1+\left( -1+\delta_{H} \right)e_{H} \right)\left( -1+f \right) \right)+ \delta_{A}\left( -1+f\left( 1+\left( -1+\delta_{H} \right)e_{H}+f-\delta_{H}f \right) \right) \right)d_{A}d_{H}^{2})$$

$$k_{3}=\left( a_{A}f^{2}I \right)\left( \frac{a_{A}e_{H}l(a_{H}e_{H}I-\left( -1+\delta_{A}+e_{H}-\delta_{H}e_{H} \right)d_{A}d_{H}}{a_{H}e_{H}l+a_{A}\left( 1+\left( -1+\delta_{H} \right)e_{H} \right)d_{H}}+\frac{a_{A}d_{H}(d_{A}+\left( -1+ \delta_{A} \right)fd_{A}+\frac{(1+\left( -1+\delta_{H} \right)f^{2})(a_{A}a_{H}e_{H}I-a_{A}\delta_{A}d_{A}d_{H})}{a_{H}e_{H}l+a_{A}\left( 1+\left( -1+\delta_{H} \right)e_{H} \right)d_{H}}}{a_{H}} \right)^{-1}$$
