## Supplemental Information 6 for "Integrating ecosystem and contaminant models to predict the effects of ecosystem fluxes on contaminant dynamics"

**Appendix S6:** Future directions for fully incorporating dynamic feedbacks between biomass and contaminants

We used a simple ecosystem coupled contaminant model to demonstrate how ecosystem processes can influence contaminant dynamics which in turn can feedback and influence ecosystem processes. These chemicals, however, are often designed with specific purposes in mind, for example acute, deleterious effects on insects deemed pest species by agriculture (e.g., neonicotinoid pesticides; Gill, Ramos-Rodriguez, and Raine 2012), but can have adverse effects at even higher trophic levels (e.g., the decline of peregrine falcons; Hellou, Lebeuf, and Rudi 2013). While the focus of our work was to use a relatively simple framework and demonstrate how spatial ecosystem processes, such as increasing nutrient inputs or spatial fluxes from donor ecosystem, can influence contaminant dynamics, this framework can be modified to explore more complex questions. These more complex questions include how those elevated contaminant concentrations can, in turn, alter food web processes such as increasing mortality rate (Pašková et al. 2011), impairing development (Mañosa et al. 2001, Story and Cox 2001), or changing behaviour (Fry 1995). One study examining effects of pesticide runoff on aquatic communities demonstrated that species known to be sensitive to pollutants had significantly lower abundances as concentration of pesticide in runoff increased even though the environmental concentration of contaminant that these species were being exposed to were far below (< 1/100) of their *LC_50_* (or the concentration of contaminant required to kill 50 % of the organisms, a traditional and frequently measured endpoint in ecotoxicology — Liess and Ohe 2005; see Table 1 in Fleeger, Carman, and Nisbet 2003 for more examples of direct and indirect effects of contaminants on ecosystems).

One way we can explore the deleterious impacts of these chemicals on ecosystems is by modifying the recycling rate of trophic level *i* (*d_i_*; which includes mortality, egestion, and excretion). Specifically, we can separate this term into a mortality rate (*m_i_*) and an egestion and excretion rate, or abscission rate for autotrophs (*g_i_*). Modelling a contaminant with a direct impact on mortality (e.g., the pesticides presented by Fleeger, Carman, and Nisbet 2003), we can then change mortality rate from being a fixed parameter to one which is a function of the contaminant concentration in that trophic level. That is:

$$m_{i}=f(\frac{C_{i}}{A, S,or H})$$

Since the early 1990s there has been a steady increase in studies in exploring the impact of specific contaminants on particular species (see Figure 1 in Köhler and Triebskorn 2013). Many of these studies explore the toxicity of a chemical using a dose-response curve; examining how a population of organisms may be affected at different levels of exposures (see Ritz 2010 for various approaches to modeling these). These curves are sigmoidal and monotonic and can be fit using a Hill Equation (Neubig et al. 2003). That is:

$$m_{i}=\frac{1}{1+\left( \frac{LC_{50}}{(\frac{C_{i}}{B_{i}})} \right)^{s_{1}}}$$

In this way, at low concentrations of contaminant in the organism the impact on mortality is low, while at higher concentrations of contaminant in the organism (*C_i_/(A, S, or H)* where *C_i_* is the biomass of trophic level *i*) the mortality rate approaches 1, with the point of inflection occurring around the *LC_50_*, and the rate of increase from zero to one being determined by *s_1_* (see Appendix S6: Figure S1). Note for the ease of this example we used the *LC_50_*, however, any other effect concentration including a lowest observed effect concentration could be used instead.

When contaminant effects on mortality are incorporated into our model we see that qualitative relationships are conserved, but small quantitative difference emerge (see Appendix S4: Figure S5 & S6). For example, adding a feedback of contaminant concentration on mortality rate decreases the equilibrium biomass of each trophic level which translates into an average higher equilibrium concentration of contaminant in that trophic level, particularly in the herbivore (*H*). Moreover, the more positive the *s_1_* exponent is, the higher the equilibrium contaminant concentration (see Appendix S4: Figure S5 & S6). This is just one way that contaminant feedback effects can be incorporated into our framework, and even within this simple example, contaminant effects on mortality may differ by trophic level or species specific sensitivity to the contaminant (e.g., Cetinić, Previšić, and Rožman 2021). Other effects of contaminants include disruption to behaviour. For example, there is evidence that exposure to neonicotinoid pesticides impacts bumblebee foraging behaviour, learning, and memory (Belzunces et al. 2012, Blacquiere et al. 2012, Farooqui 2013). By adapting consumption rate (parameter *a_i_* in our model) our framework can be used to explore these scenarios.

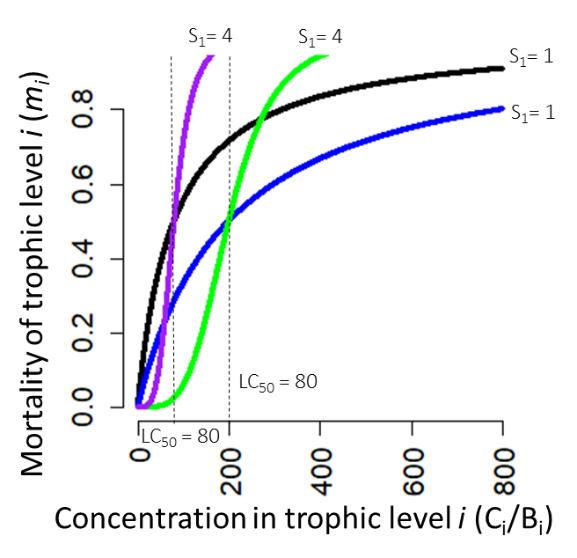

**Figure S1.** Relationship between mortality of trophic level *i* and the concentration of contaminant in that trophic level. Lines indicate two different values of the exponent *s_1_*, i.e., *s_1_* = 4 (purple and green), and *s_1_* = 1 (blue and black); and two different LC_50_ values, i.e., LC_50_ = 80 (purple and black) and LC_50_ = 200 (blue and green) where values for *s_i_* would be obtained from best literature estimates of lethal concentrations. Alternatively, this could be modeled as a more conventional dose-response curve, where instead of mortality as a function of the concentration in trophic level *i* (*C_i_/*$(A, S, or H)$), we consider the total exposure for trophic level *i*. In this way, $m_{i}=\frac{1}{1+\left( \frac{LC_{50}}{(w_{i}u_{i}E_{0}+u_{i}(A, S, or H)E_{1}+fa_{i}(A, S, or H)C_{i-1})/(A, S, or H)} \right)^{s_{1}}}$.

**References:**

Belzunces, L. P., S. Tchamitchian, and J.-L. Brunet. 2012. Neural effects of insecticides in the honey bee. Apidologie 43:348–370.

Blacquiere, T., G. Smagghe, C. A. Van Gestel, and V. Mommaerts. 2012. Neonicotinoids in bees: a review on concentrations, side-effects and risk assessment. Ecotoxicology 21:973–992.

Cetinić, K. A., A. Previšić, and M. Rožman. 2021. Holo-and hemimetabolism of aquatic insects: Implications for a differential cross-ecosystem flux of metals. Environmental Pollution 277:116798.

Farooqui, T. 2013. A potential link among biogenic amines-based pesticides, learning and memory, and colony collapse disorder: a unique hypothesis. Neurochemistry international 62:122–136.

Fleeger, J. W., K. R. Carman, and R. M. Nisbet. 2003. Indirect effects of contaminants in aquatic ecosystems. Science of The Total Environment 317:207–233.

Fry, D. 1995. Reproductive effects in birds exposed to pesticides and industrial chemicals. Environmental Health Perspectives 103:165–171.

Gill, R. J., O. Ramos-Rodriguez, and N. E. Raine. 2012. Combined pesticide exposure severely affects individual-and colony-level traits in bees. Nature 491:105–108.

Hellou, J., M. Lebeuf, and M. Rudi. 2013. Review on DDT and metabolites in birds and mammals of aquatic ecosystems. Environmental Reviews 21:53–69.

Köhler, H.-R., and R. Triebskorn. 2013. Wildlife ecotoxicology of pesticides: can we track effects to the population level and beyond? Science 341:759–765.

Liess, M., and P. C. V. D. Ohe. 2005. Analyzing effects of pesticides on invertebrate communities in streams. Environmental Toxicology and Chemistry 24:954–965.

Mañosa, S., R. Mateo, and R. Guitart. 2001. A review of the effects of agricultural and industrial contamination on the Ebro delta biota and wildlife. Environmental Monitoring and Assessment 71:187–205.

Neubig, R. R., M. Spedding, T. Kenakin, and A. Christopoulos. 2003. International Union of Pharmacology Committee on Receptor Nomenclature and Drug Classification. XXXVIII. Update on terms and symbols in quantitative pharmacology. Pharmacological reviews 55:597–606.

Pašková, V., K. Hilscherová, and L. Bláha. 2011. Teratogenicity and embryotoxicity in aquatic organisms after pesticide exposure and the role of oxidative stress. Reviews of Environmental Contamination and Toxicology 211:25–61.

Ritz, C. 2010. Toward a unified approach to dose–response modeling in ecotoxicology. Environmental Toxicology and Chemistry 29:220–229.

Story, P., and M. Cox. 2001. Review of the effects of organophosphorus and carbamate insecticides on vertebrates. Are there implications for locust management in Australia? Wildlife Research 28:179–193.
